# Asymmetry-induced transient gel formation in fluid lipid membranes

**DOI:** 10.1101/2025.09.11.674739

**Authors:** Emad Pirhadi, Xin Yong

## Abstract

Compositional asymmetry is a defining feature of cellular membranes, controlling permeability, protein activity, cholesterol dynamics, and shape remodeling. This asymmetry can create a stress imbalance, with the two leaflets experiencing opposing tensions, though direct experimental measurement of leaflet stress remains challenging. Such a stress imbalance can compress one leaflet and trigger a fluid-to-gel phase transition, which reduces membrane fluidity and markedly increases bending rigidity. These phenomena raise a key question of how membranes respond mechanically before crossing the transition threshold, a regime that remains relevant to biological functions. Here, we combine comprehensive all-atom and coarse-grained molecular dynamics simulations to examine how stress asymmetry modulates membrane structure and mechanics near the transition point. Using POPE and DLPC bilayers as model systems, we find that moderate asymmetry induces transient gel-like domains that continuously form and dissolve, amplifying undulations and lowering bilayer rigidity. Beyond the gelation threshold, the trend reverses and the bilayer stiffens, resulting in a non-monotonic dependence of rigidity on asymmetry. Moreover, our results reveal distinct curvature preferences of fluid and gel phases. Extending this analysis to a multicomponent bacterial outer membrane, we demonstrate that stress asymmetry can trigger transient gel-like domain formation even in complex lipid mixtures. This provides a proof of principle that differential stress modulates membrane mechanics by inducing either softening or stiffening, complementing the effects of molecular composition. Our findings elucidate how cells might exploit the stress-curvature-phase coupling to tune membrane rigidity under near-physiological conditions.

**Statement of significance:** Cell membranes are selectively permeable bilayers composed of different lipids that are unevenly distributed both across the two leaflets and within each leaflet. These structural features influence how membranes maintain their integrity, position proteins, and send signals. While each feature has been studied separately, the direct influence of leaflet asymmetry on lateral lipid organization remains unclear. Using molecular simulations, we show that leaflet asymmetry can drive lipids to cluster into coexisting fluid and gel domains at body temperature. This reorganization softens the membrane and enhances its fluctuations. Our findings reveal a direct link between leaflet asymmetry and lateral structure, shedding light on how cells can tune membrane properties to regulate communication, transport, and adaptation to stress.

## Introduction

Cellular membranes function as dynamic, selective barriers that maintain homeostasis while supporting diverse processes such as signaling, trafficking, and mechanical deformation.(1–3) These essential roles are closely linked to two defining biophysical features: compositional asymmetry between the two leaflets and lateral heterogeneity within each leaflet.(4–7) The asymmetric distribution of lipids is essential for numerous cellular functions, including vesicle formation, membrane trafficking, and signal transduction.(8–11) By creating differential leaflet stress, asymmetry can drive or suppress curvature generation, regulate membrane protein orientation and activity, and modulate the recruitment of curvature-sensing proteins.(12–16) In parallel, lateral heterogeneity organizes membrane components into dynamic functional domains that further influence protein activity.(17–22) Laterally, lipid bilayers can adopt a phase-separated state, with liquid-ordered (Lo) domains characterized by tightly packed and more ordered lipids coexisting with liquid-disordered (Ld) regions that are more fluid and loosely packed.(23) Lateral phase separation in bilayers is known to depend on lipid headgroup chemistry, tail length and saturation, charge, and the presence of cholesterol.(23–25) Cholesterol, in particular, plays a dual role by stabilizing both ordered and disordered phases depending on the surrounding lipid environment.(26–28) Moreover, lipid phase behavior is highly sensitive to temperature; below the main phase transition temperature (*T_m_*), the bilayer can transition from a fluid phase to a gel phase, characterized by reduced lateral mobility and increased stiffness.(29–31)

While the thermotropic behavior of membranes has been extensively studied, recent findings suggest that stress asymmetry can also influence phase behavior. Coarse-grained simulations by Hossein and Deserno (32,33) demonstrated that stress asymmetry can drive phase transitions, even at constant temperature. Depending on proximity to *T_m_*, such stress can induce fluid-gel coexistence and alter bilayer mechanical properties.(34) Importantly, while both Lo and gel (Lβ) phases involve high tail ordering, they differ fundamentally in lateral dynamics: Lo domains are fluid and cholesterol-rich, whereas gel domains are solid-like and typically cholesterol-poor.(35) From a physical standpoint, the gel phase exhibits unique and intriguing properties, however, its biological relevance remains debated as extreme rigidity can compromise membrane function and cell viability.(36–38) In both eukaryotic and prokaryotic cells, lipid compositions are tightly regulated to maintain a *T_m_* slightly above the culture temperature.(39,40) This raises the intriguing possibility that cells exploit stress asymmetry to tune membrane behavior near the phase transition boundary, maintaining global fluidity while enabling transient formation of ordered domains.

In this study, we employ comprehensive all-atom and coarse-grained molecular dynamics (MD) simulations to investigate how stress asymmetry, introduced via leaflet lipid mismatch, modulates phase behavior and mechanical properties in both simplified and physiologically relevant membrane models. We find that, near the phase transition limit, bilayers remain globally fluid yet form transient gel-like domains that display curvature sensitivity, enhanced undulations, and reduced bending rigidity at a constant physiologically relevant temperature. Furthermore, under large differential stress, our all-atom simulations confirm recent observations of membrane stiffening and gel-phase formation in coarse-grained studies.(32–34) This reveals a two-staged mechanical response of the membrane to asymmetry: softening near the transition threshold, followed by gelation-induced stiffening. Together, our results suggest a mechanistic framework by which cells tune their membrane deformability and lateral organization via leaflet asymmetry. This mechanism may enable adaptive responses to environmental cues, regulate protein activity, or enhance resistance to membrane-targeting agents such as antimicrobial peptides. By linking compositional asymmetry to dynamic control over phase behavior, this study provides new insights into how both eukaryotic and prokaryotic cells may regulate membrane mechanics at physiological temperatures.

## Methods

### All-atom MD simulations

All-atom simulations were conducted using the CHARMM36 force field.(41,42) 1- palmitoyl-2-oleoyl-sn-glycero-3-phosphoethanolamine (POPE) bilayers contained only POPE lipids and water. Lipid A bilayers were composed of hexa-acylated *Pseudomonas aeruginosa* (*P. aeruginosa*) Lipid A (PA14 strain), with calcium ions added for charge neutralization. The bacterial outer membrane (OM) model was constructed based on experimental lipidomics data,(43–45) with the outer leaflet consisting of Lipid A and the inner leaflet containing a 2.2:1 mixture of POPE and 1-palmitoyl-2-oleoyl-sn-glycero-3-phosphoglycerol (POPG). Sodium and calcium ions were introduced to neutralize the charges of POPG and Lipid A, respectively, and 150 mM NaCl was added to mimic physiological conditions. NBFIX (46,47) corrections were applied to improve the accuracy of ion-lipid and ion-ion interactions.

Initial configurations were generated using the CHARMM-GUI membrane builder,(48) and simulations were carried out with GROMACS 2023.1.(49,50) Each system underwent energy minimization via the steepest descent algorithm, followed by 100 ps of initial equilibration under an NVT ensemble using the velocity-rescaling thermostat,(51) with separate coupling for lipids and solvent. Final equilibration and production simulations were then performed in the NPT ensemble using semi-isotropic pressure coupling at 1 bar. Although semi-isotropic coupling is standard for fluid lipid membranes, the emergence of gel phases and gel-like domains warrants verification of its suitability. To assess the robustness of this choice, selected systems (one in the stable gel phase and one in the pre-transition regime) were also simulated using anisotropic NPT pressure coupling with independent control over the *x*, *y*, and *z* box dimensions. Structural and mechanical properties obtained from these simulations were in close agreement with those from semi-isotropic simulations, and the equilibrated box dimensions in the *x* and *y* directions were comparable after extended sampling (see Fig. S1 and Table S1). These results support the appropriateness of semi-isotropic coupling for the systems studied here. Additional details are provided in the Supplemental Information (SI). All reported production data in the main manuscript were generated using the stochastic cell rescaling barostat (52) and stochastic velocity rescaling thermostat.(53) Equilibration durations ranged from 2 to 5 μs. All bonds involving hydrogen atoms were constrained using the LINCS algorithm.(54) Short-range van der Waals and electrostatic interactions were truncated at 1.2 nm, with neighbor lists updated every 20 steps. Long-range electrostatics were treated using the Particle-Mesh Ewald (PME) method.(55) A 2 fs integration timestep was used with the leapfrog integrator. Detailed simulation parameters, system compositions, and data sampling information are provided in SI.

### Coarse-grained MD simulations

Coarse-grained simulations were performed using the MARTINI 2.0 force field.(56) A timestep of 20 fs was employed. Following established protocols, Lennard-Jones and Coulomb interactions were truncated at 1.1 nm, and a relative dielectric constant of 15 was used. Thermostats and barostats were implemented using stochastic velocity rescaling and stochastic cell rescaling, respectively. Neighbor list parameters were set according to the recommendations in Ref. (57).

### Hidden Markov Model

Gaussian Hidden Markov Model (GHMM) was used to assign lipid phase states based on per-lipid structural features.(58) A two-state GHMM was implemented to represent ordered and disordered phases. Features included area per lipid, carbon-carbon order parameter, and tail height. Initial emission distributions were estimated by fitting a Gaussian Mixture Model (GMM) to a system exhibiting clear phase separation. The GHMM was then trained using the Viterbi algorithm.(59) To ensure robustness against local minima, the model was trained under 10 different random seeds, and the one with the highest log-likelihood was selected. GMM and GHMM analyses were performed using the scikit-learn (60) and hmmlearn Python packages, respectively.

### Bending modulus

We employed three independent methods to estimate the bending modulus (𝜅) from structural fluctuations: (1) height fluctuations in Fourier space (hq),(61,62) (2) transverse curvature bias (TCB) analysis,(63) and (3) real-space fluctuations (RSF) based on lipid splay.(64,65)

1. **Fourier-space height fluctuations**

The bending modulus was estimated by analyzing the power spectrum of membrane height fluctuations, following the Helfrich continuum elastic theory.(66) The membrane surface was discretized using a grid with a spacing of approximately 15 Å, and height values were computed using atoms located at the neutral surface, defined as the surface where the transverse curvature bias vanishes (see next section). The Fourier components of the surface height were calculated using:

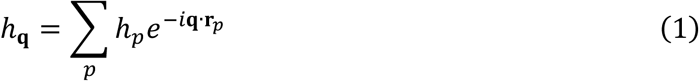

The fluctuation spectrum fits the following expression:

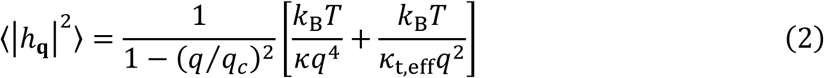

where ℎ_𝐪_ is the Fourier amplitude of mode 𝐪, 𝐫_𝑝_ and ℎ_𝑝_ are the position and height of grid point 𝑝, 𝜅_t,eff_ is the effective tilt modulus, and 𝑞_𝑐_ is the critical wavenumber fit parameter. This approach captures both the standard Helfrich 𝑞^4^ scaling and corrections due to tilt contributions. See Ref. (62) for additional information on the theory.

2. **Transverse curvature bias**

The TCB method measures the curvature experienced by lipid molecules at various heights *z* above or below the membrane midplane. It relies on the coupling between lipid director and local membrane curvature. The curvature sampled by lipids at a height *z* from the neutral surface is related to the bending modulus by:

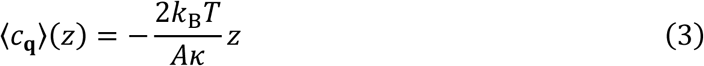

where 〈𝑐_𝐪_〉(𝑧) is the mean curvature of mode 𝐪𝐪 at height *z* away from the neutral surface and 𝐴 is the membrane area. A linear fit of curvature versus height yields the bending modulus (Fig. S2).

3. **Real-space fluctuations**

The RSF method correlates the bending modulus to fluctuations in lipid splay, defined as the deviation between the lipid director vector (𝐧𝐧) and the normal vector (N) to the water-membrane interface. The local splay is given by:

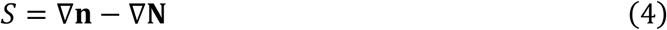

Assuming a Boltzmann distribution for the splay, the bending modulus is extracted from the distribution of splay values:

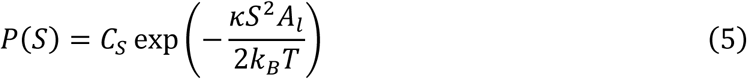

where 𝐴_𝑙_ is the area per lipid, 𝑃(S) is the probability distribution of splays, and 𝐶_𝑆_ is the normalization constant. This method captures local curvature information at the level of lipid pairs and is particularly suited for analyzing homogeneous or fluid systems with short-range ordering.(65)

In the presence of fluid-gel coexistence, the membrane is laterally heterogeneous, and the Helfrich assumption of a uniform material no longer strictly applies. The 𝜅 values reported here should therefore be interpreted as effective bending moduli, representing the large-scale bending response of a composite membrane whose local mechanical properties vary spatially.(33)

## Results

### Leaflet asymmetry drives a fluid-to-gel phase transition

We investigated how leaflet asymmetry influences bilayer lateral heterogeneity using all-atom and coarse-grained molecular dynamics simulations. Asymmetry was introduced by creating a lipid number mismatch between the two leaflets, quantified as

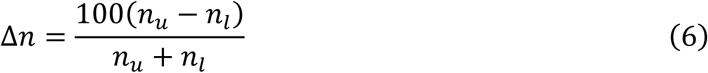

where 𝑛_𝑢_ and 𝑛_𝑙_ are the numbers of lipids in the upper and lower leaflets, respectively. Δ𝑛 represents the percentage difference in lipid count between leaflets and is directly related to differential stress (see Table S2): overpopulating one leaflet imposes negative tension (compression) on that leaflet and positive tension (extension) on the opposite leaflet. Leaflet number asymmetry is used here as a surrogate for differential stress, rather than as a direct representation of physiological lipid distributions. This approach isolates the mechanical consequences of interleaflet stress imbalance on membrane structure. Simulations were performed with POPE (Fig. 1A,B), chosen for the significant presence of PE in plasma membranes (particularly the cytoplasmic leaflet) and its established role in membrane budding and cell division.(67–69) POPE lies close to its phase transition at physiological temperature,(70) making it particularly sensitive to compressive stress and therefore well suited for detecting stress driven structural responses. Because the ordering observed here is leaflet-specific, the mechanism does not rely on POPE being distributed across both leaflets and remains relevant to asymmetric biological membranes where PE is enriched in the inner leaflet.(71) Nonetheless, the generality of the observed behavior across lipid chemistry, force fields, and realistic multicomponent systems is examined in subsequent sections. Bilayer phase separation has been shown to be size dependent, thus we tested three system sizes containing approximately 200, 400, and 800 lipids per leaflet (Table S3). Fig. S3 shows that phase separation behavior is consistent among these system sizes, indicating that the observed effects are not artifacts of system size within the length scales accessible here. The rest of our analysis on POPE bilayers is done on the ∼400 lipid/leaflet set.

**Figure 1.**
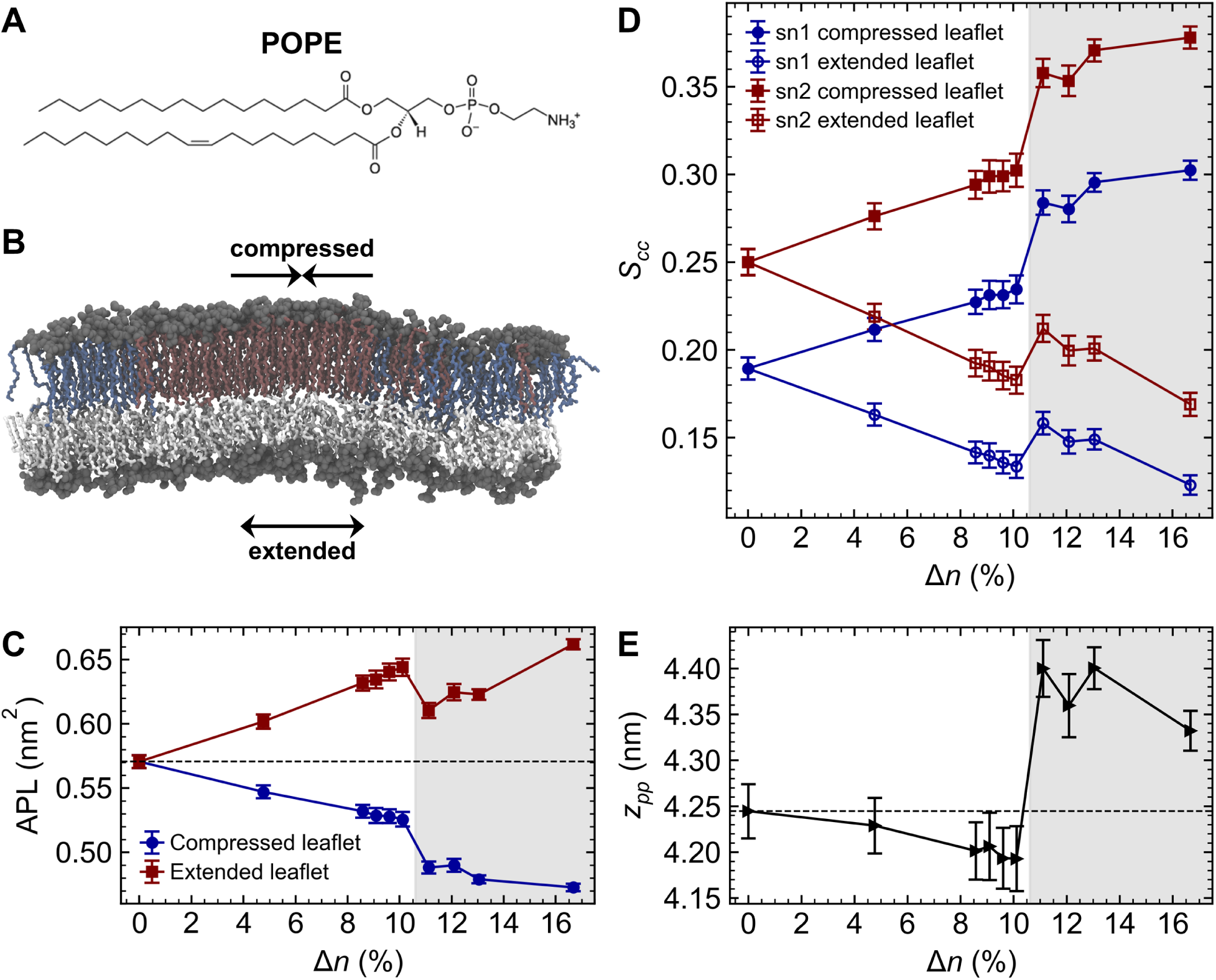
Stress asymmetry modulates bilayer structure. (A) Chemical structure of POPE. (B) Representative simulation snapshot of a POPE bilayer under number asymmetry (Δ𝑛 ≈ 13.0%). Lipid tails in the extended leaflet are shown in white, while lipid tails in the compressed leaflet are colored by phase: gel phase (dark red) and fluid phase (blue). (C) Area per lipid of each leaflet, (D) carbon-carbon order parameter per leaflet, and (E) phosphate-to-phosphate membrane thickness as functions of number asymmetry. Shaded regions indicate the post phase transition regime. Error bars represent standard deviations of the mean over time.

As shown in Fig. 1C, increasing Δ𝑛 leads to a sharp decrease in the average area per lipid (APL) in the compressed leaflet at Δ𝑛 ≈ 11.1%, consistent with enhanced lipid packing. At this same degree of asymmetry, we observed a marked increase in lipid tail carbon-carbon order parameters (𝑆_𝑐𝑐_) and bilayer thickness (𝑧_𝑝𝑝_) (Fig. 1D,E), signatures of a transition to the gel phase.(72) Beyond this threshold, the gel-phase compressed leaflet dominates overall bilayer properties, as reflected in the bilayer thickness. Notably, prior to the transition, the extended (positive tension) leaflet exhibits greater sensitivity to asymmetry. This is evident from the more pronounced deviation in APL and 𝑆_𝑐𝑐_ of the extended leaflet before the phase transition (Fig. 1C,D). As a result, bilayer thickness decreases by approximately 0.05 nm due to an overall less ordered bilayer within this limit. These changes indicate that differential stress affects bilayer properties even in the absence of a phase transition.

### Ordered and disordered domains display opposite curvature preferences in a fluid-gel mixture

Above the asymmetry threshold Δ𝑛 ≥ 11.1%, the compressed leaflet exhibits a mixture of ordered gel domains and disordered fluid regions, consistent with the observations in coarse-grained simulations of 1,2-dilauroyl-sn-glycero-3-phosphocholine (DLPC) bilayers reported by Foley et al.(34) We mapped the instantaneous structural properties of individual lipids in the compressed leaflet and visualized their spatial distribution as heatmaps (see Fig. 2A-C). The distinct degree of lipid packing visible in these figures, together with quantitative differences in tail order parameter, tail height, and area per lipid, confirms the presence of two structurally distinct domains. This coexistence remains stable throughout our analysis simulation time (2 µs), with small fluctuations in the gel domain size (see Videos S1-3). In contrast, the extended leaflet remains homogeneous and does not exhibit gel domain formation within this range of number asymmetry (see Fig. S4).

**Figure 2.**
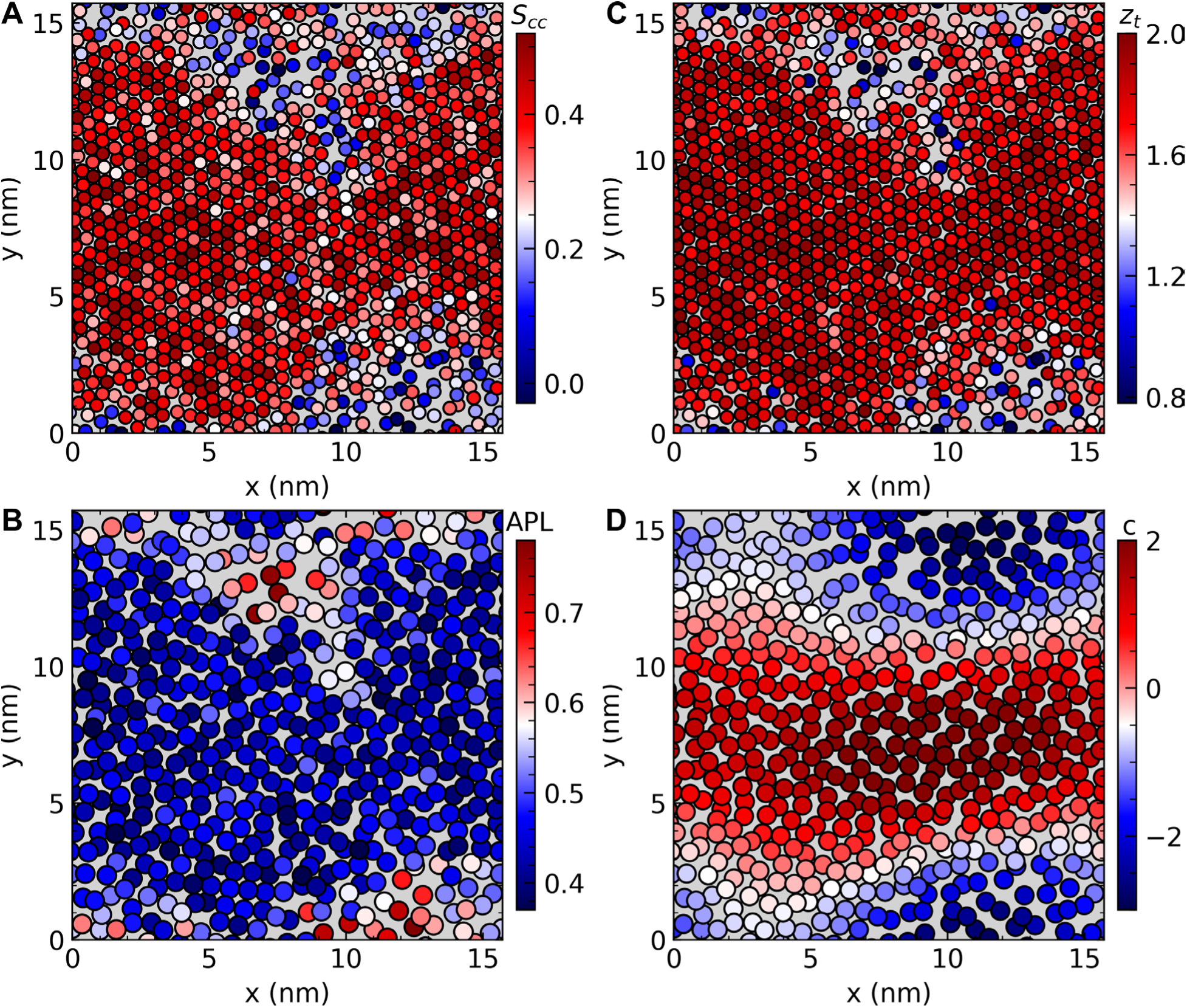
Stress asymmetry induces lateral phase separation. Heatmaps of structural properties for the compressed leaflet of a POPE membrane under number asymmetry of Δ𝑛 ≈ 13.0%, plotted for the final simulation snapshot: (A) carbon-carbon order parameter, (B) area per lipid, (C) tail height in the z-direction, and (D) curvature experienced by each lipid. Each bead corresponds to (A,C) the center of mass of an individual lipid tail or (B,D) the average center of mass of both tails.

To examine curvature sensitivity, we computed the instantaneous curvature experienced by each lipid from the Fourier spectrum of bilayer height fluctuations(63,73):

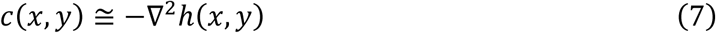

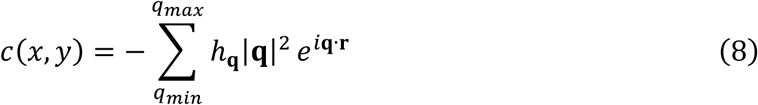

where ℎ_𝐪_ is the Fourier amplitude of mode 𝐪, 𝐫_𝑝_ and ℎ_𝑝_ are the position and height of grid point 𝑝, 𝑞_𝑚𝑖𝑛_ and 𝑞_𝑚𝑎𝑥_ define the range of 𝐪 used in the calculation, and 𝑐(𝑥, 𝑦) is the local curvature. We focused on low 𝐪 modes (𝑞_𝑚𝑎𝑥_ = 0.8 nm^−1^) to capture large-scale undulations while avoiding significant noise in larger 𝐪𝐪 modes. Representative snapshots (see Fig. 2D) show that gel domains are consistently registered with regions of positive curvature, whereas fluid regions occupy areas of negative curvature. This curvature partitioning persists throughout the entire 2 µs of our analysis trajectory.

To quantify structural differences between phases, we apply a Hidden Markov Model (HMM),(58) which uses an unsupervised machine learning algorithm to classify each lipid as gel or fluid.(74,75) We train the model using three per-lipid attributes: tail order parameter, tail height, and area per lipid. Assuming a binary phase state, the HMM labels each lipid in every frame, enabling phase-specific curvature statistics. As shown in Fig. 3, the curvature difference between gel and fluid domains increases with asymmetry and exhibits a distinct jump at the phase transition threshold (Δ𝑛 ≥ 11.1%). These findings reveal that asymmetry-induced phase coexistence is accompanied by distinct curvature preferences of fluid and gel domains.

**Figure 3.**
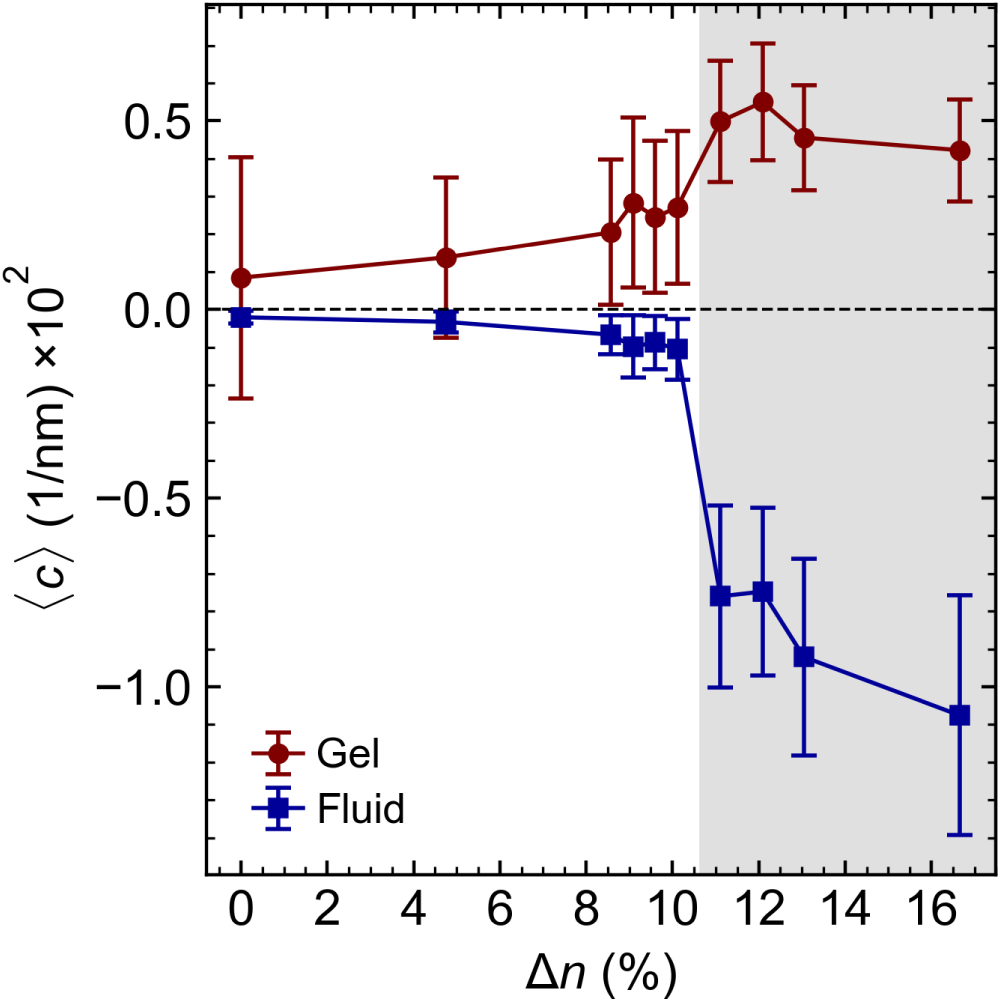
Lipid phase state modulates curvature preference. Time-averaged local curvature experienced by fluid and gel phase lipids in the compressed leaflet of POPE membranes, plotted as a function of number asymmetry. The shaded region marks the post phase transition regime. Error bars represent standard deviations of the mean over time.

### Transient domain formation drives bilayer softening near the asymmetry-induced phase transition

The gel fraction in the compressed leaflet increases gradually with Δ𝑛 until the large-scale phase transition (Δ𝑛 ≈ 11.1%), where it exhibits a sharp jump (Fig. 4A). Beyond this point, gel domain growth is attenuated, and 100% gel fraction would likely require asymmetry levels that compromise bilayer stability. Herein, we focus on the pre-transition regime 8.6% ≤ Δ𝑛 ≤ 10.1%, where the bilayer remains globally fluid, which is a physiologically plausible state.

**Figure 4.**
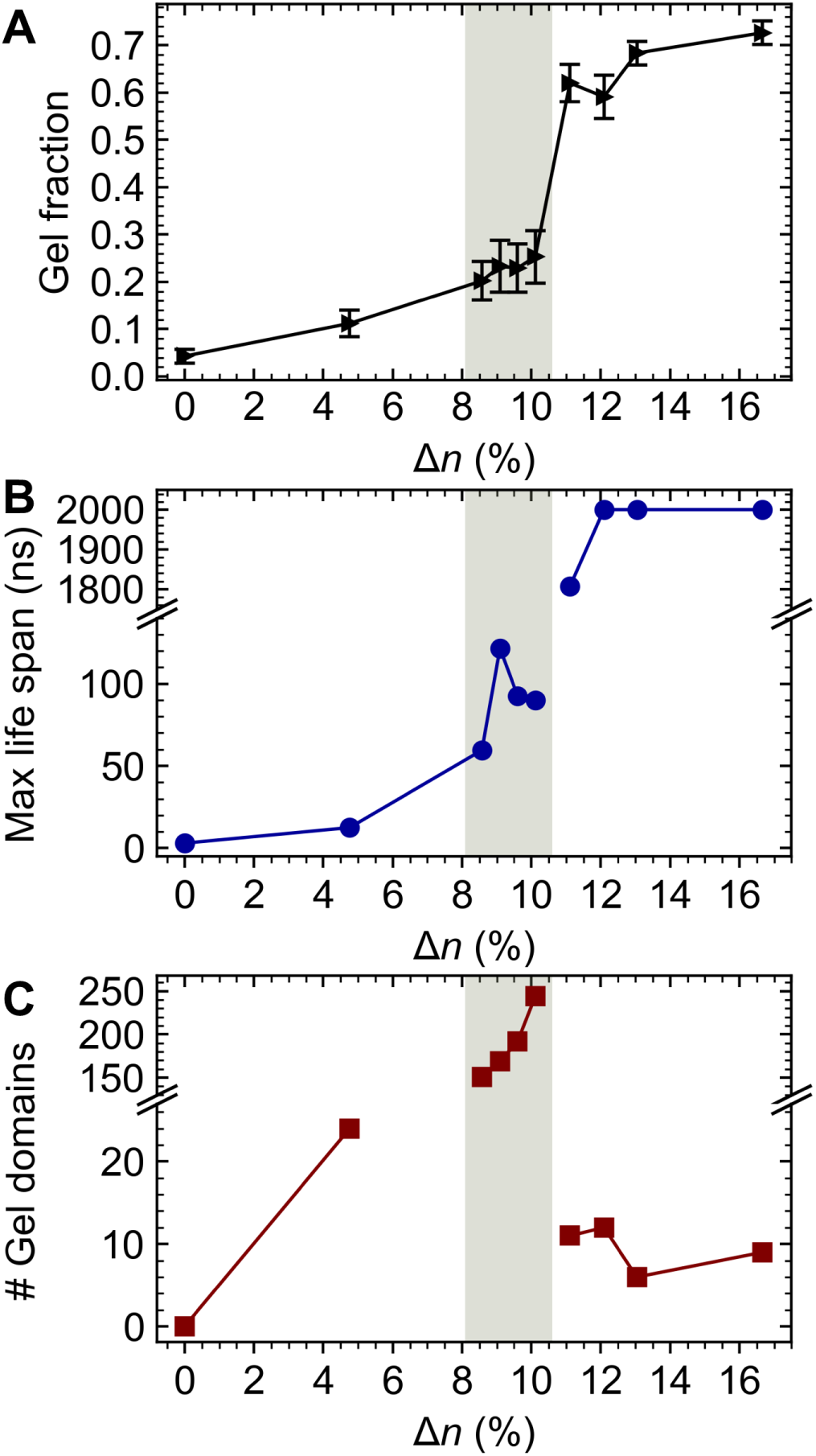
Asymmetry promotes transient gel-like domain formation near the transition threshold. (A) Fraction of gel-phase lipids in the compressed leaflet of POPE membranes, (B) maximal lifetime of gel clusters during the 2-μs simulation, and (C) total number of gel domains formed throughout the trajectory as functions of number asymmetry. Shaded areas highlight the near-transition regime exhibiting notable transient gel-like domains. Error bars denote standard deviations of the mean over time.

HMM-based phase mapping and spatiotemporal domain analysis reveal that in this range, small gel-like domains form transiently within the fluid leaflet (see Videos S1–S3). A gel-like domain was defined as a minimum of 4 neighboring gel-state lipids persisting for at least 5 ns. The number of domains and their maximum lifetime both increase with asymmetry until the global transition, at which point separate domains merge into a single stable one during the entire span of simulation. In the pre-transition regime, more than 200 domains form during the entire trajectory, with maximal lifetimes greater than 100 ns. The combination of this clustering behavior and a gel fraction of 20-30% (Fig. 4B,C) indicates highly dynamic phase heterogeneity without complete solidification in the compressed leaflet.

Previous work by Hossein and Deserno (32) reported an asymmetry-induced increase in the bilayer bending modulus following a fluid-to-gel phase transition, consistent with thermodynamic expectations for comparing homogeneous gel and fluid phases. In contrast, our analysis targets the pre-transition regime, where transient domains appear. Here, we find evidence of softening rather than stiffening in that regime.

We quantified mechanical properties by computing the area compressibility modulus (𝐾_𝐴_) and effective bending modulus (𝜅) as functions of number asymmetry (Δ𝑛). The area compressibility modulus was calculated from equilibrium fluctuations in bilayer area under the zero membrane tension condition.(76) The decrease in 𝐾_𝐴_ with increasing asymmetry in the pre-transition regime indicates bilayer softening, which is followed by a sharp increase upon gelation (Fig. 5B). For 𝜅, we applied three standard methods, height fluctuations in the Fourier space (hq),(61,62) transverse-curvature-bias (TCB),(63) and real-space fluctuations (RSF).(64,65) Unfortunately, all three methods rely on assumptions valid only in the fluid phase, in which the membrane cannot sustain in-plane shear stress and the Helfrich Hamiltonian is applicable.(66) In the gel state, these assumptions are violated, and the fluid membrane Helfrich description is no longer appropriate. This can be seen by the power spectra of undulations that illustrate a pronounced q-dependence in the gel state (see Fig. S5). A continuum elastic model suitable for the solid-like nature of gel phase is required to describe this regime. The RSF method shows a monotonic increase in 𝜅 with increasing asymmetry, even before gelation. However, this apparent stiffening may be an artifact as RSF is largely insensitive to large-scale fluctuations and instead reflects local lipid splay dominated by short-range interactions. As a result, while RSF is a useful tool for homogeneous membranes, it likely reports local lipid ordering rather than the true bending elasticity in our heterogeneous systems (see Fig. S6). Consistent with this limitation, a recent study has reported that RSF overestimates the bending modulus in cholesterol-rich membranes,(77) suggesting that it may fail to capture diffusional softening mechanisms.(78) To the extent of our knowledge, the buckling method (79,80) often used for coarse-grained models may be the only method capable of calculating the bending modulus in the gel phase. However, this method assumes a mirror-symmetric buckle geometry, which follows from a homogeneous continuum elastic description. In asymmetric membranes exhibiting phase coexistence, gel domains can preferentially localize at one inflection point, systematically breaking this symmetry.(33) This makes the predicted buckle shape only approximate, although still providing a meaningful estimate of the effective bending modulus. Additionally, because the buckling method imposes curvature, it could inadvertently promote phase transitions in membranes near *T_m_*, especially if curvature favors the gel phase over the fluid phase (see Fig. 3). This may explain the apparent pre-transition stiffening reported by Ref. (32).

**Figure 5.**
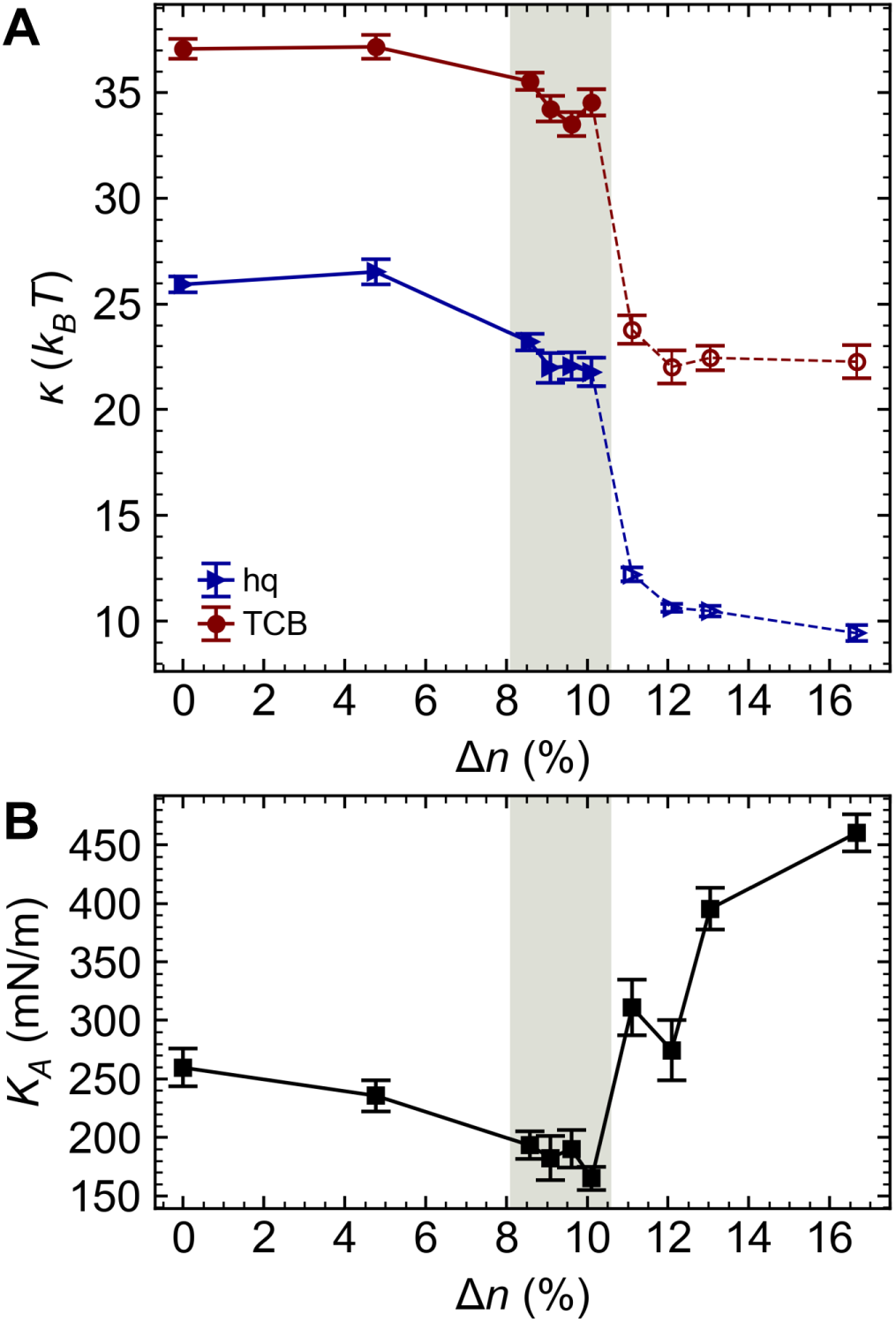
Membrane softening near the phase transition threshold. (A) Bending modulus and (B) area compressibility modulus plotted as functions of number asymmetry. Error bars represent standard errors of the mean obtained via block averaging (see SI for details). Shaded areas highlight the near-transition regime which exhibits notable transient gel-like domains.

The hq method, which is considered the gold standard for fluid bilayers, shows a clear and statistically significant decrease in 𝜅 with increasing asymmetry in the range 8.6% ≤ Δ𝑛 ≤ 10.1% (Fig. 5A and S7). This reduction in bending resistance is consistent with the behavior of 𝐾_𝐴_, confirming bilayer softening near the phase transition. A similar trend is observed using the TCB method, though the absolute values of 𝜅 are shifted. The observed pre-transition softening resembles the “anomalous swelling” phenomenon reported in temperature-driven transitions by several experimental studies.(81–83)

The presence of curvature-sensitive inclusions in fluid membranes is known to induce softening, as described by the classic theory of Leibler.(84) This softening mechanism attributed to the coupling of curvature to membrane undulations has been supported by both computational and experimental studies.(78,85–87) Here, we tested whether this framework can account for the softening observed in our simulations by approximating transient gel-like domains as a single curvature-sensitive inclusion of effective area 𝐴_𝐺_ within an otherwise fluid bilayer.(88) A prefactor of 1 ⁄ 2 is applied to account for an inclusion in a single leaflet. Under this assumption, the expected reduction in bending rigidity can be estimated as

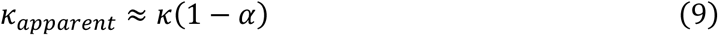

with

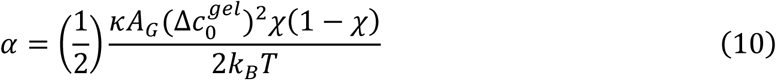

Here, 𝜒 is the mole fraction of gel-like domains, 𝐴_𝐺_ is the effective area of the inclusion, and 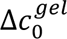 is the spontaneous curvature difference relative to the fluid background. For Δ𝑛 = 10.1%, we obtain 𝜒 = 0.22, 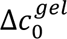 = 0.012 nm^−1^, and 𝐴 = 57.9 nm^2^. Using the symmetric bilayer bending modulus 𝜅 = 25.5 𝑘_𝐵_𝑇, this yields an estimated softening of only ∼1%. This prediction is, however, significantly smaller than the ∼16% softening obtained directly from the hq method (Fig. 5A). This discrepancy reflects the limitations of the inclusion theory in this regime. The theory assumes static, well-defined inclusions embedded in a homogeneous fluid membrane with negligible inclusion–inclusion interactions. In contrast, gel domains here are transient, fluctuate in size and number, interact, and create a mechanically heterogeneous membrane. These effects likely introduce additional long-wavelength fluctuations that are not captured by the static inclusion model. Moreover, the theoretical prediction is highly sensitive to Δ𝑐^𝑔𝑔𝑒𝑙^, so small uncertainties in curvature sampling can lead to large deviations in the predicted softening. We therefore conclude that the inclusion-based softening theory does not fully account for the softening observed here. Instead, the additional reduction in apparent bending rigidity arises from enhanced bilayer dynamics associated with the continual formation and dissolution of gel-like domains in the pre-transition regime.

### Asymmetry-induced phase behavior depends on lipid chemistry and force field

We assessed the generality of asymmetry-induced phase behavior by simulating additional systems, including coarse-grained (CG) DLPC bilayers using MARTINI 2.0, and all-atom Lipid A bilayers (Tables S4-5). In all-atom Lipid A at 310 K, no phase transition occurred before reaching the bilayer’s stress tolerance limit, beyond which bilayer integrity failed as the compressed leaflet expelled a lipid (Fig. S8). Further simulations showed a low *T_m_* of ∼290 K for Lipid A, making it unlikely that stress asymmetry would induce gel formation at physiological temperature (Fig. S9). By contrast, CG DLPC in MARTINI 2.0 did exhibit a fluid-to-gel phase transition when simulated at 300 K, close to its reported *T_m_*.(89) Consistent changes in tail order parameter, area per lipid, and bilayer thickness (see Fig. 6A-C) indicated a transition at Δ𝑛 ≈ 14.0% (Fig. S10). This range is considerably higher than the ≈ 6% reported phase transition limit by Ref. (32). The shift is attributable to the slightly elevated simulation temperature (+2.5 K) of this study and also to the shorter 1.1 nm cutoff for Lennard-Jones and Coulomb interactions recommended in recent MARTINI protocols.

**Figure 6.**
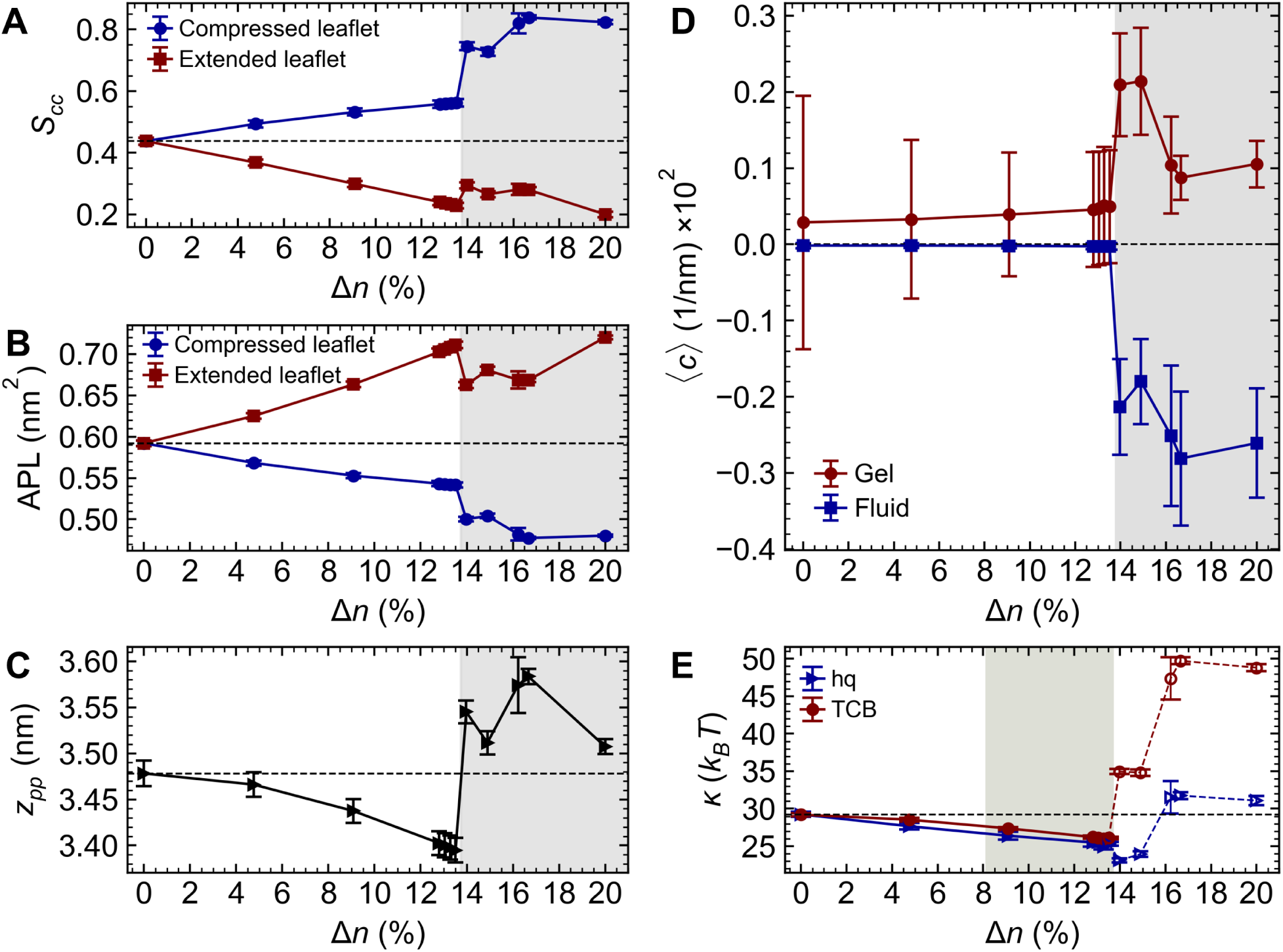
Softening near the transition regime in coarse-grained DLPC membranes. (A) Carbon-carbon order parameter, (B) area per lipid, (C) phosphate-to-phosphate membrane thickness, (D) time-averaged local curvature experienced by fluid and gel phase lipids, and (E) bending modulus as functions of number asymmetry. For panels A– D, error bars represent standard deviations over time; shaded regions indicate the onset of phase transition. In panel E, error bars show the standard error of the mean from block averaging; the shaded region denotes the near-transition regime.

For DLPC, 𝐾_𝐴_ increases sharply after gel formation (Fig. S11), consistent with the stiffening reported by Ref.(32) Moreover, both area compressibility and bending moduli show a softening trend near the transition threshold, accompanied by the formation of transient gel-like domains, consistent with our all-atom POPE results (Fig. 6D,E and S9). The apparent stiffening observed in the bending modulus after gelation is expected thermodynamically but not quantitatively reliable, as computational methods to extract the bending modulus become impractical in the gel state (Fig. S5).

### Extension to complex bacterial outer membrane systems

To evaluate asymmetry-driven structural responses beyond glycerophospholipid membranes, we simulated the outer membrane (OM) of *Pseudomonas aeruginosa* bacteria using all-atom molecular dynamics. The OM was composed of PA14 Lipid A in the outer leaflet and a mixture of POPE and POPG phospholipids (PLs) in the inner leaflet (Table S6). This system incorporates multiple relevant complexities, including compositional asymmetry, a multicomponent inner leaflet, charged lipids, and divalent cations that form ion bridges between Lipid A phosphate groups, making it a physiologically relevant testbed for our hypothesis.

Due to compositional asymmetry, the direction of asymmetry is important in this system. Asymmetry was parameterized relative to the membrane with minimum differential stress (0LT), generated using the zero-leaflet-tension method. We define asymmetry as:

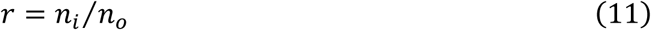

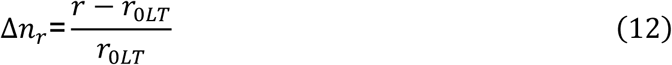

where 𝑛_𝑜_ and 𝑛_𝑖_ are the number of lipids in the outer and inner leaflets, respectively, and 𝑟_0𝐿𝑇_ is the ratio in the zero differential stress state.(90) Positive Δ𝑛_𝑟_ corresponds to inner leaflet compression (PLs leaflet), while negative Δ𝑛_𝑟_ corresponds to outer leaflet compression (Lipid A leaflet). Table S7 shows a monotonic correlation between Δ𝑛_𝑟_ and differential stress.

Consistent with our single-component results, compressing the outer leaflet (Lipid A) did not induce a phase transition, and this unique leaflet resisted compression until membrane integrity failed. In contrast, compressing the phospholipid inner leaflet (Δ𝑛_𝑟_ > +25.0%) led to the formation of ordered domains within the multicomponent PL leaflet. As shown in Fig. 7A-C, domain formation was accompanied by increased tail order parameter, decreased area per lipid, and locally high lipid packing (see Video S4). Similar to the simpler membranes, these ordered domains displayed curvature sensitivity, consistently residing in regions of positive curvature (Fig. 7D). These results demonstrate that asymmetry-driven domain formation and curvature preference, first observed in single-component bilayers, extend to multicomponent bacterial outer membranes, reinforcing the potential physiological relevance of this mechanism.

**Figure 7.**
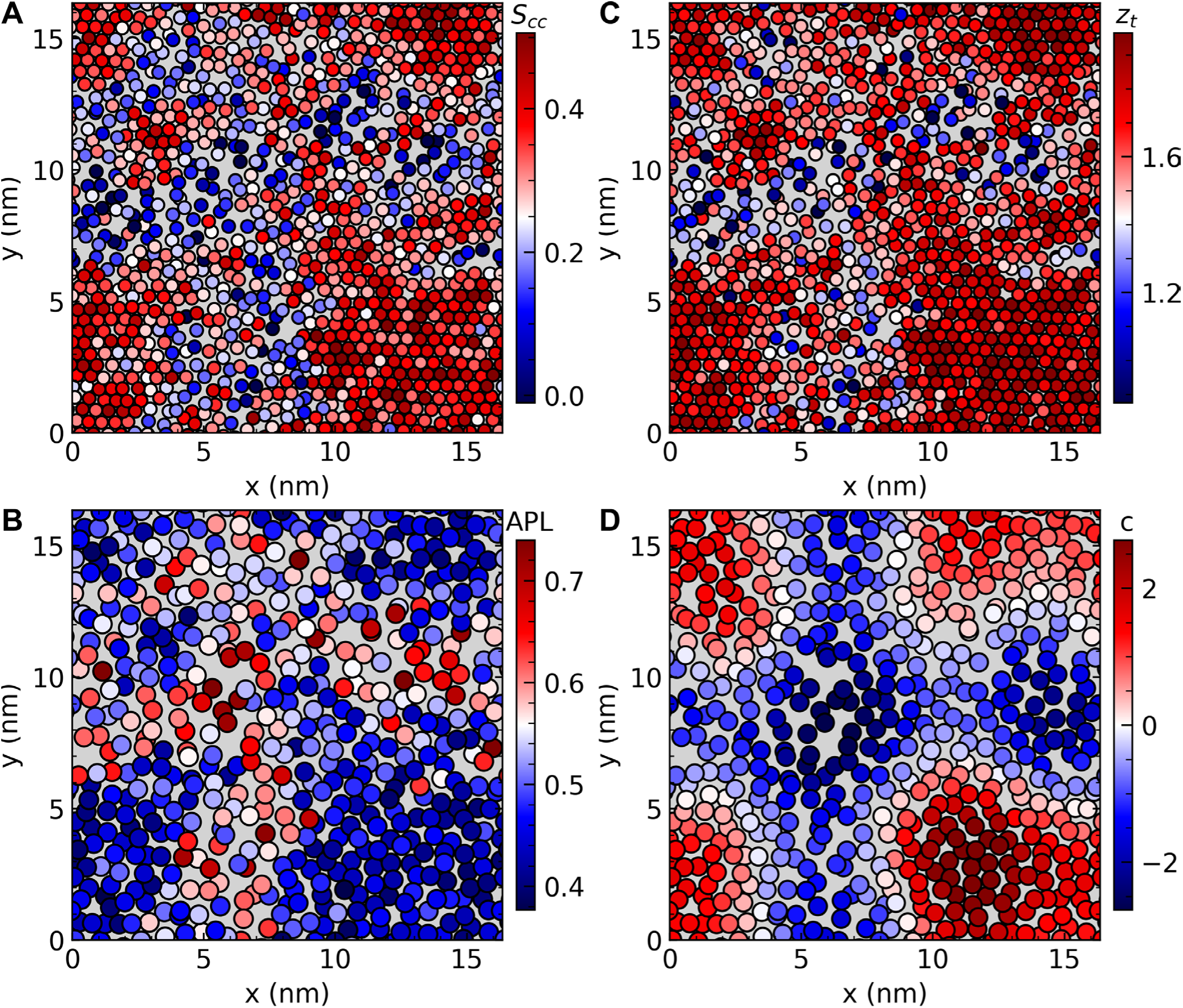
Stress asymmetry drives phase transition in the PLs leaflet of the OM. Heatmaps of structural properties for the compressed (PLs) leaflet under relative number asymmetry of Δ𝑛_𝑟_ ≈ 30.0%, plotted for the last simulation snapshot: (A) carbon-carbon order parameter, (B) area per lipid, (C) tail height in the z-direction, and (D) curvature experienced by each lipid. Each bead corresponds to (A,C) the center of mass of an individual lipid tail or (B,D) the average center of mass of both tails.

## Discussion and conclusion

Since the pioneering works of Bretscher (91) and Verkleij et al. (92) in the 1970s, membrane asymmetry has been recognized as a defining feature of biological membranes. Cells employ specialized transporter proteins known as flippases, floppases, and scramblases to control lipid asymmetry by actively translocating lipids between the two leaflets.(9,93) Leaflet asymmetry can also arise from asymmetrically shaped transmembrane proteins, leaflet-specific lipid domains, or the anchoring of the cytoplasmic leaflet to submembrane networks such as the cytoskeleton or peptidoglycan layer.(11,94) In addition to these compositional effects, cell membranes may also maintain number asymmetry, where the two leaflets contain unequal quantities of identical lipids or molecules.(12,71) This number asymmetry alters lipid packing and induces leaflet-specific strain, resulting in differential stress, a condition in which lateral tension differs between the two leaflets. Although differential stress is relatively easy to observe and control in computational modeling,(32,90) direct experimental quantification remains challenging due to limitations in measuring microscopic stress in bilayers.(95) However, its effects can be inferred indirectly. For example, asymmetric giant unilamellar vesicles (GUVs) are remarkably stable despite their low curvature, which is two orders of magnitude lower than the spontaneous curvature of their component lipids. This implies the presence of an opposing mechanical contribution, which may arise from differential stress balancing the bending torque. This likely differential stress has been estimated to be in the order of several mN/m.(96) Additionally, differential stress has been predicted to influence several biological functions, including lipid flip-flop, membrane budding, activation of transmembrane proteins, peptide-induced pore formation, and the intercalation of small molecules.(12,13,15,90,97,98)

Recent studies have shown that differential bilayer stress can actively promote phase transitions and enhance lateral heterogeneity.(32,33) Near the main phase transition temperature *T_m_*, compressive stress can drive lipid molecules into the gel state. Depending on the magnitude of the stress, these gel domains may coexist with fluid regions, forming a stable heterogeneous mixture.(34) In this regime, number asymmetry is kinetically conserved on the simulation timescale and therefore acts as an additional intensive thermodynamic control variable. As a result, two phase coexistence can occupy a finite region rather than a single line in the phase diagram.(34) The connection between mechanical stress and phase behavior suggests a regulatory mechanism by which cells can modulate membrane fluidity and permeability, thereby broadening the physiological significance of lipid asymmetry. In addition to mechanical stress, several other factors including temperature, cholesterol content, ion concentration, and membrane-associated proteins contribute to the modulation of membrane phase behavior.(17,26,30,99) Notably, the spontaneous curvature of lipids has been hypothesized to correlate with lateral organization and the overall phase state of membranes.(100,101) Supporting this idea, experiments on asymmetric POPE/POPC large unilamellar vesicles (LUVs) (102) have shown that POPE, which has a *T_m_* of ∼25 °C, undergoes a phase transition at 16 °C when placed in the inner leaflet, but at 20 °C when located in the outer leaflet. This upward shift is attributed to curvature-induced differential stress as POPE with negative spontaneous curvature is forced to adopt positive curvature when positioned in the outer leaflet. These observations underscore the role of curvature-induced stress in modulating phase behavior, a connection supported by our results. For the systems examined in this study, we find that within a given leaflet, gel domains preferentially localize to regions of positive curvature, while disordered fluid regions are enriched in negatively curved areas (Fig. 3 and 6D). This spatial preference is consistent with the increase in spontaneous curvature driven by the tight packing and ordering of hydrocarbon chains. Overall, the observed phase distribution illustrates a dynamic and reciprocal interplay between membrane curvature and lipid phase state. Such curvature-phase coupling may also influence, or be influenced by, the localization and activity of curvature-sensing proteins, providing an additional feedback mechanism linking membrane structure to function.

Several independent experiments have reported higher bending rigidity in asymmetrical vesicles compared to their symmetrical counterparts.(103–106) Coarse-grained simulations have attributed this stiffening to differential stress,(32,33,107) which can push bilayers toward the gel phase when near *T_m_*. Interestingly, a recent experiment reported asymmetry-induced stiffening even in the absence of notable differential stress, suggesting additional complexity.(103) It is important to recognize the limitations of current experimental techniques. Accurately determining leaflet asymmetry remains challenging, and the physical effects of chemical probes used in mechanical measurements are not yet fully understood.(108) Despite these uncertainties, the collective evidence clearly supports that asymmetry substantially influences membrane mechanical properties.

From a broader perspective, while gel phases exhibit rich physical properties and are commonly observed *in vitro* and *in silico*,(33,34,72,109) their suppression of bilayer fluidity renders them largely incompatible with *in vivo* conditions, where dynamic membrane behavior is essential for cell function.(36,37,110) Our simulation snapshots display a dramatic loss of fluidity upon a fluid-to-gel phase transition, in agreement with previous literature.(111) Therefore, we argue that a more physiologically relevant regime lies just below the transition threshold, where the membrane is under differential stress, near *T_m_*, but maintains a globally fluid state. A recent computational study reported a modest (<25%) increase in bending rigidity under asymmetry, even in the absence of visible lateral heterogeneity.(112) These systems were treated as homogeneous throughout the simulation. The explanation offered centered on the greater sensitivity of van der Waals forces to compression than extension, resulting in the compressed leaflet dominating bilayer mechanics. However, in our simulations, we do not observe statistically significant stiffening within the low asymmetry regime where the bilayer remains fluid.

Intriguingly, as the system approaches the transition threshold, we observe a reduction in both the area compressibility modulus and effective bending modulus, indicating a non-monotonic mechanical response to increasing asymmetry. This softened regime is characterized by enhanced lateral fluctuations, driven by the transient formation of ordered gel-like domains. These domains increase area oscillation, thereby reducing area compressibility modulus. Concurrently, curvature analysis reveals that gel domains localize in highly curved areas (Fig. 3 and 6D). The spontaneous appearance and disappearance of gel-like domains locally modulate curvature, contributing to dynamic feedback that amplifies undulatory fluctuations. A similar curvature instability on a global scale has been observed in coarse-grained Monte Carlo simulations under conditions of phase coexistence.(107) The sensitivity of transient gel-like domains to curvature may be a key contributor to the softening effect we observe. Their formation promotes localized bending, and their eventual dissolution relaxes curvature. This mechanism resonates with recent results from Ref. (113), where lipid mixtures with contrasting spontaneous curvatures induced membrane softening through nonadditive structural effects.(114,115) However, curvature-composition coupling theory (84,88) underestimates the degree of softening observed in our simulations. We speculate that the transient nature of gel-like domain formation introduces an additional dynamic mode of softening, whereby time-dependent recruitment and relaxation of curved domains amplifies membrane undulations. This interpretation aligns with Schick’s theoretical framework,(116) in which curvature-composition coupling gives rise to transient “microemulsion” domains that reduce the bending energy of bilayer fluctuations, even in the absence of full phase separation. However, the underlying mechanisms of softening are likely more complex. Collective effects, such as inter-leaflet coupling, may further modulate membrane mechanics in the presence of asymmetric transient domains. Targeted investigations will be necessary to disentangle these contributions and clarify the origins of the softening observed in this regime. Finally, it is worth noting that enhanced bilayer fluctuations near phase transitions have also been observed experimentally.(38,117) The reduction of bending rigidity in this regime is frequently associated with the phenomenon of anomalous swelling.(81–83)

One may question whether the observed softening near the transition is simply a consequence of bilayer thinning. While the polymer-brush model may not hold in this regime, we emphasize that the observed ∼16% decrease in bending modulus and 36% decrease in compressibility modulus cannot be explained by the ∼2% reduction in thickness. We note that in the vicinity of phase transition, enhanced fluctuations in elastic properties are expected as a hallmark of critical behavior. However, the softening trends observed here are consistent across independent systems. Another potential critique concerns the transient ordered domains “flakes” reported here, and whether they are simply part of natural fluctuations. However, our analysis shows that these domains can reach ∼130 lipids in size and persist for over 100 ns, significantly larger and longer-lived than background fluctuations in symmetric bilayers (see Fig. 4B and S12B). One caveat of our study is the difficulty of sampling near phase transition regimes using all-atom force fields. Transient domain formation slows relaxation and increases autocorrelation times, requiring extended trajectories. Although we performed 2 μs equilibration followed by 2 μs analysis, longer simulations would likely provide more definitive evidence for the observed behavior. Additionally, it is important to note that the elastic moduli reported here represent global averages over the entire bilayer. This is a standard approach when analyzing fluid membranes. However, with the onset of gel-like domain formation, the mechanical properties of individual domains and leaflets will inevitably vary. Even so, the apparent average moduli remain informative and relevant to membrane shape remodeling.

To assess the generality of these findings, we extended our analysis to a bacterial outer membrane model that better reflects physiological complexity. Despite this system’s increased compositional and structural complexities, we still observed a similar fluid-to-gel phase transition. This model, of course, introduces additional complications and requires further study to be fully understood. For instance, the presence of divalent ion bridges in the outer leaflet (composed of Lipid A) likely increases its tolerance to negative differential stress, which results from crowding and compression of the inner PL leaflet. We acknowledge that including the full lipopolysaccharide structure instead of only the Lipid A anchor could potentially alter the observed behavior. However, the added sugars would likely increase the number of ion bridges, and thus could further enhance stress tolerance. Regardless, the key takeaway from this complex model is the plausibility of accessing the near-transition regime in a physiologically relevant context.

Our findings support and extend previous studies on the impact of asymmetry on membrane structure and function.(13,32–34,106) We propose that both eukaryotic and prokaryotic cells may use asymmetry as a regulatory mechanism to fine-tune their lateral phase behavior. This modulation could support various biological functions, including protein recruitment and activation, fluidity control, and defense against environmental stress. For instance, bacteria might exploit asymmetry-induced stiffening to resist antimicrobial peptides that disrupt membrane mechanics,(118) or induce softening to facilitate vesiculation.(119,120)

## Author contributions

X. Y. conceived and supervised the research. E. P. designed and performed the MD simulations, conducted the analyses, and prepared the figures. E. P. wrote the manuscript, and X. Y. contributed to editing and revision. Both authors discussed the results and approved the final version of the manuscript.

## Acknowledgments

We thank Alexander Sodt for assistance in MembraneAnalysis.jl package, constructive discussion, and critical reading of the manuscript. This work was supported in part by U.S. National Science Foundation grant #2448213 and National Institutes of Health grant 1R15GM135862. Computational resources of this research were provided by the Center for Computational Research at the University at Buffalo and allocation BIO220055 from the Advanced Cyberinfrastructure Coordination Ecosystem: Services & Support (ACCESS) program, which is supported by U.S. National Science Foundation grants #2138259, #2138286, #2138307, #2137603, and #2138296.

## Supporting citations

References (121–129) appear in the supplemental information.

## Pressure coupling scheme comparison

Semi-isotropic and anisotropic pressure coupling schemes were compared to assess the robustness of the semi-isotropic coupling used in this work. Because the stochastic cell-rescaling barostat (1) implemented in GROMACS does not support fully anisotropic pressure coupling, anisotropic test simulations were performed using the Parrinello-Rahman barostat.(2) For the anisotropic simulations, the *x*, *y*, and *z* box dimensions were coupled independently, while shear terms (off-diagonal components of the compressibility matrix) were disabled to prevent box shear deformation. This choice avoids unphysical shear distortions, particularly given that portions of the membrane remain in the fluid phase and the surrounding solvent does not sustain shear.

Two representative POPE all-atom systems were examined: one in the stable gel phase (Δ𝑛 = 13.0%) and one in the pre-transition regime (Δ𝑛 = 9.6%). Table S1 confirms that the average box dimensions under different pressure coupling schemes are similar. As shown in Fig. S1, structural properties of the membrane with stable fluid-gel coexistence, obtained under anisotropic coupling, are comparable to those from semi-isotropic simulations. Transient gel domain formation at Δ𝑛 = 9.6% was also consistently observed under both coupling schemes. These results indicate that the main structural conclusions are not sensitive to the choice of pressure coupling scheme within the regimes examined.

Second-order pressure coupling schemes such as Parrinello-Rahman often require a longer pressure relaxation time constant than first-order schemes (e.g., stochastic cell rescaling) to ensure numerical stability and avoid box oscillations.(3) Accordingly, 𝜏_𝑝_ = 10 ps was used for Parrinello-Rahman, compared to 𝜏_𝑝_ = 5 ps for stochastic cell rescaling. The longer relaxation time contributes to the increased fluctuation amplitudes reported in Table S1. Importantly, the qualitative structural behavior and phase characteristics remain unchanged.

## Structural properties

### 1. Order parameter

To quantify lipid tail ordering, we calculated both the carbon-hydrogen order parameter (𝑆_𝑐ℎ_) and the carbon-carbon order parameter (𝑆_𝑐𝑐_).(4) These metrics assess the orientational order of lipid tails relative to the membrane normal. The 𝑆_𝑐ℎ_ parameter, appropriate for all-atom simulations, is defined based on the angle 𝜃𝜃_𝑐ℎ_ between the bilayer normal and the C–H bond vector:

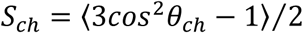

The 𝑆_𝑐𝑐_ parameter, commonly used in coarse-grained simulations, is calculated using the angle 𝜃𝜃_𝑐𝑐_ between the bilayer normal and the vector connecting consecutive tail carbons or beads:

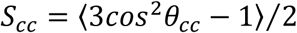

We employed LipidDyne (5) to calculate 𝑆_𝑐ℎ_ and LiPyphilic (6) for 𝑆_𝑐𝑐_. As shown in Fig. S13, there is excellent agreement between the two order parameters for the all-atom POPE membrane and both measures show similar dependence on number asymmetry. To maintain consistency across all-atom and coarse-grained models, 𝑆_𝑐𝑐_ was used as an input feature for the Hidden Markov Model (HMM) training throughout this study.

### 2. Area per lipid (APL)

A rough estimate of APL can be obtained from the simulation box size. However, for HMM classification, per-lipid APL values are required. To compute this, we used a 2D Voronoi tessellation implemented in the LiPyphilic package, applied to the tail carbons or beads of each lipid. This approach provides a more accurate representation of the local lipid environment and allows spatially resolved measurements of APL.

### 3. Thickness (𝑧_𝑝𝑝_)

Membrane thickness is defined as the phosphate-to-phosphate (P–P) distance between the two leaflets. To account for undulations and local curvature effects, we employed the FATSLiM (7) package. This method calculates the local normal to the membrane surface and estimates the thickness along that direction.

### 4. Tail height (𝑧_𝑡_)

Tail height is defined for each lipid as the maximum distance along the z-axis between the terminal tail atoms or beads. The calculation was conducted using LiPyphilic. This quantity captures fluctuations in lipid extension and provides an additional structural feature used in phase state analysis.

## Error estimation for bending and area compressibility moduli

Error bars shown in Fig. 5A,B were calculated using a block averaging approach. The simulation trajectory was divided into 15 blocks, each spanning 200 ns, which corresponds to the maximum lifetime of transient gel-like clusters (see Fig. 4B). For the TCB method, block values of the bending modulus were combined using the harmonic mean (i.e., averaging the inverse moduli and then inverting), and the associated error bars were obtained by propagating the uncertainties accordingly. For all other methods, block values were averaged using the arithmetic mean, and the standard error of the mean was reported as the error bar.

## Per-lipid curvature analysis over time

To quantify curvature preferences of lipids in different phase states, we used the MembraneAnalysis.jl (8) package to compute membrane curvature at the per-lipid, per-frame resolution. The original implementation was extended by modifying the source code to output local curvature values for each lipid in every frame of the trajectory. We focused on low 𝐪𝐪 modes (𝑞_𝑚𝑎𝑥_ = 0.8 nm^−1^) to capture large-scale undulations while avoiding significant noise in larger 𝐪𝐪 modes. This high-resolution curvature dataset was then combined with the HMM-assigned phase states to separately evaluate the curvature environments that lipids preferentially occupy in the gel and fluid phases. For each time frame, lipids were grouped based on their assigned states, and their corresponding curvature values were averaged. This allowed us to characterize phase-specific curvature localization and assess dynamic curvature-phase coupling over the course of the simulation.

## Transient domain characterization

To identify stable domains of ordered lipids, we defined a cluster as a group of at least four neighboring lipids that were all assigned to the gel phase by the HMM and remained in that state for a minimum of 10 consecutive simulation frames (see Fig. S14). Clustering was based on per-frame lipid coordinates and the corresponding per-lipid phase state. Neighbor search was performed using a 7-8 Å cutoff based on the estimated lipid radius from the APL. A cluster was considered persistent across frames if at least 60% of its constituent lipids were retained between successive frames. If this criterion was not met, the cluster was considered dismantled and no longer tracked.

## Local stress tensor calculation

Local stress tensor (𝛔𝛔) components and profiles were computed using GROMACS-LS,(9,10) a modified version of GROMACS that implements the central force decomposition (CFD) (11) method for calculating stress. Based on recent benchmarks,(12,13) a Coulomb cutoff of 2.8 nm was applied to accurately capture long-range electrostatic contributions to the local stress. To ensure adequate convergence of stress profiles, 400,000 frames were sampled from 2 μs of simulation data. The stress profile calculations were performed on systems containing approximately 100 lipids per leaflet to minimize fluctuations that could otherwise smear stress profiles.

Differential stress (Δ𝛾𝛾) is calculated as the difference between the tensions of outer and inner leaflets as follows:

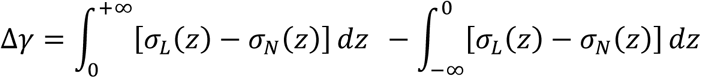

where 𝜎_𝐿_ = (𝜎_𝑥𝑥_ + 𝜎_𝑦𝑦_)/2 is the lateral stress, 𝜎_𝑁_ = 𝜎_𝑧𝑧_ is the normal stress, and the bilayer midplane is located at *z* = 0.

## Supplemental Figures

**Figure S1.**
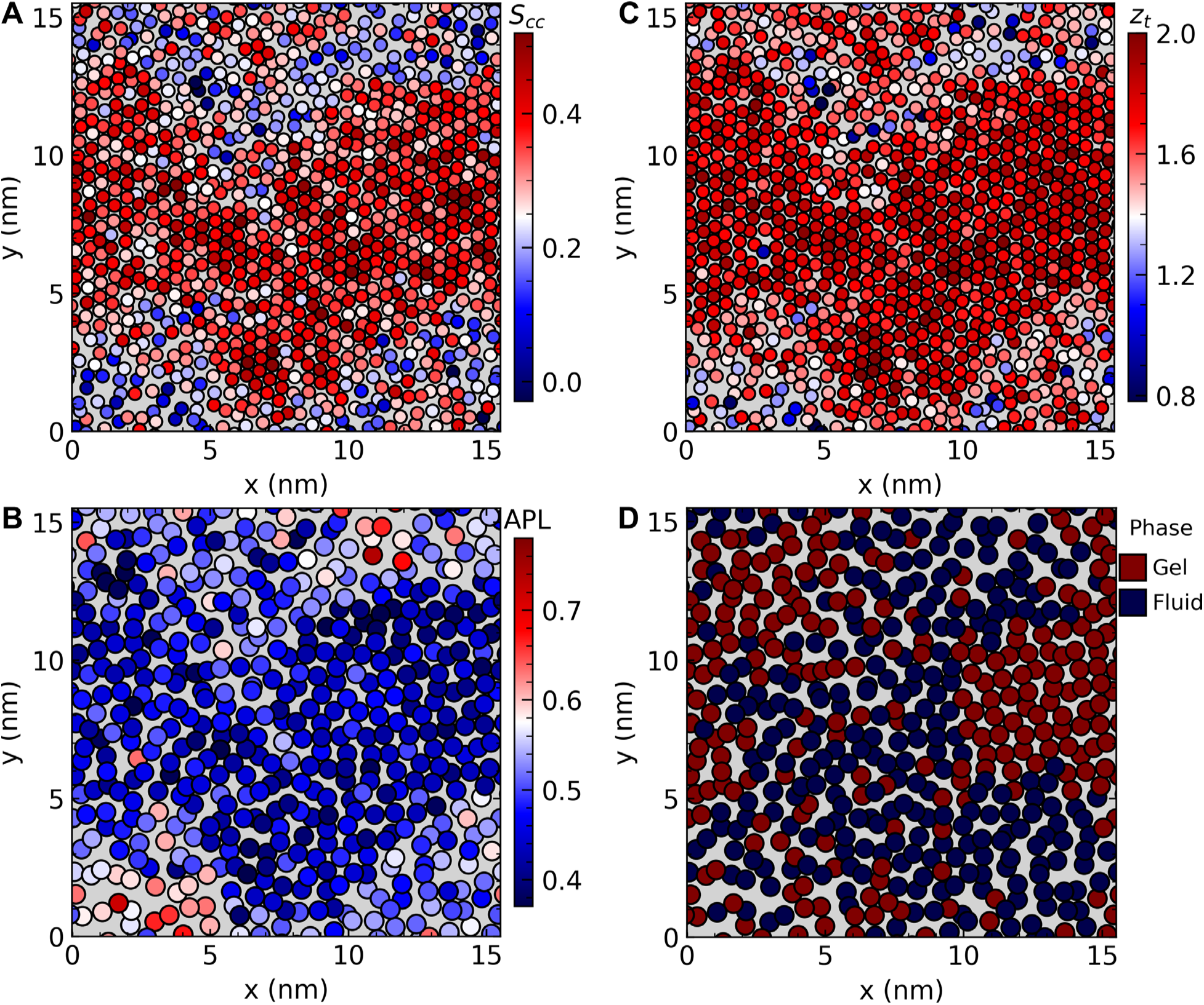
Stress asymmetry induces lateral phase separation regardless of pressure coupling scheme. Heatmaps of structural properties for the compressed leaflet of a POPE membrane under number asymmetry of Δ𝑛 ≈ 13.0%, plotted for the final simulation snapshot under anisotropic pressure coupling: (A) carbon-carbon order parameter, (B) area per lipid, (C) tail height in the z-direction, and (D) HMM-resolved phase state. Each bead corresponds to (A,C) the center of mass of an individual lipid tail or (B,D) the average center of mass of both tails.

**Figure S2.**
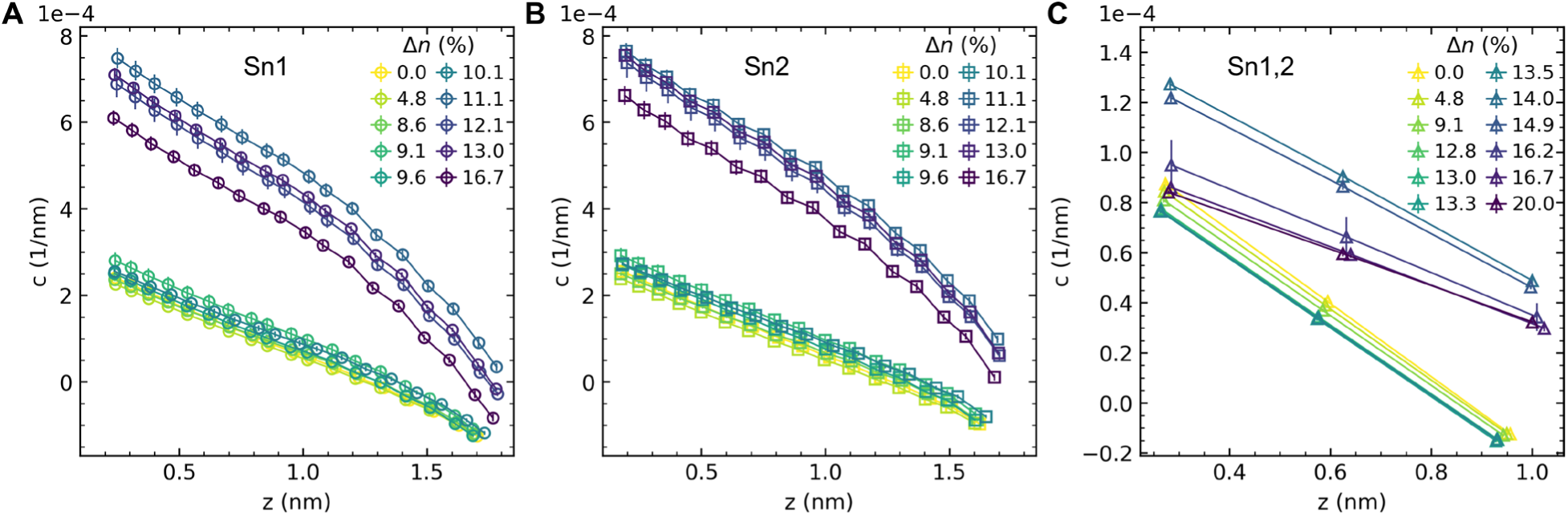
Transverse curvature bias as a function of distance from the midplane. (A) all-atom POPE sn-1 chain, (B) all-atom POPE sn-2 chain, and (C) average of both chains in MARTINI DLPC membranes.

**Figure S3.**
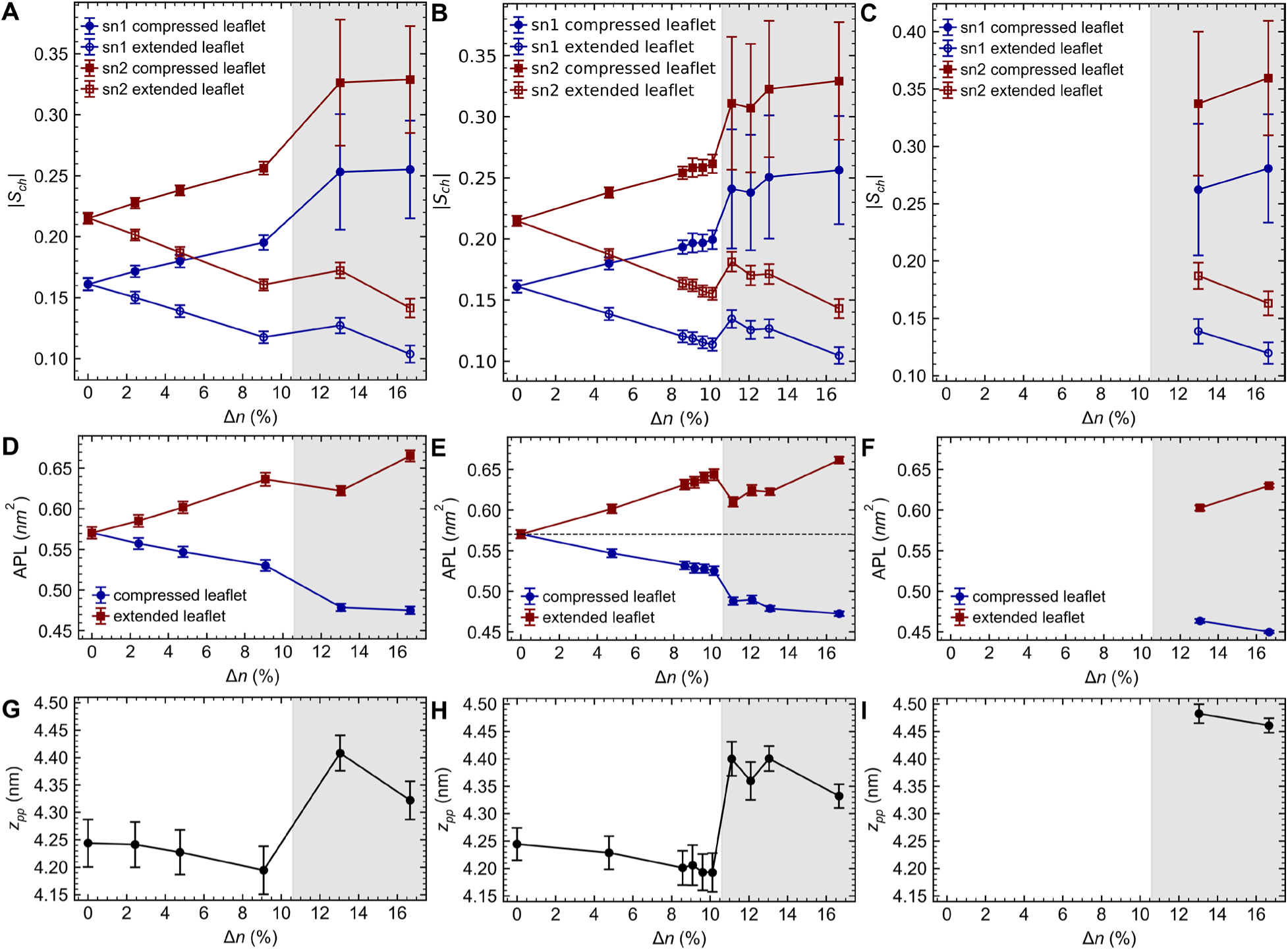
Structural properties of all-atom POPE membranes of varying sizes. (A-C) Per-leaflet carbon-hydrogen order parameter, (D-F) per-leaflet area per lipid, and (G-I) phosphate-to-phosphate membrane thickness as functions of number asymmetry. The first, second, and third columns correspond to membranes with approximately 200, 400, and 800 lipids per leaflet, respectively.

**Figure S4.**
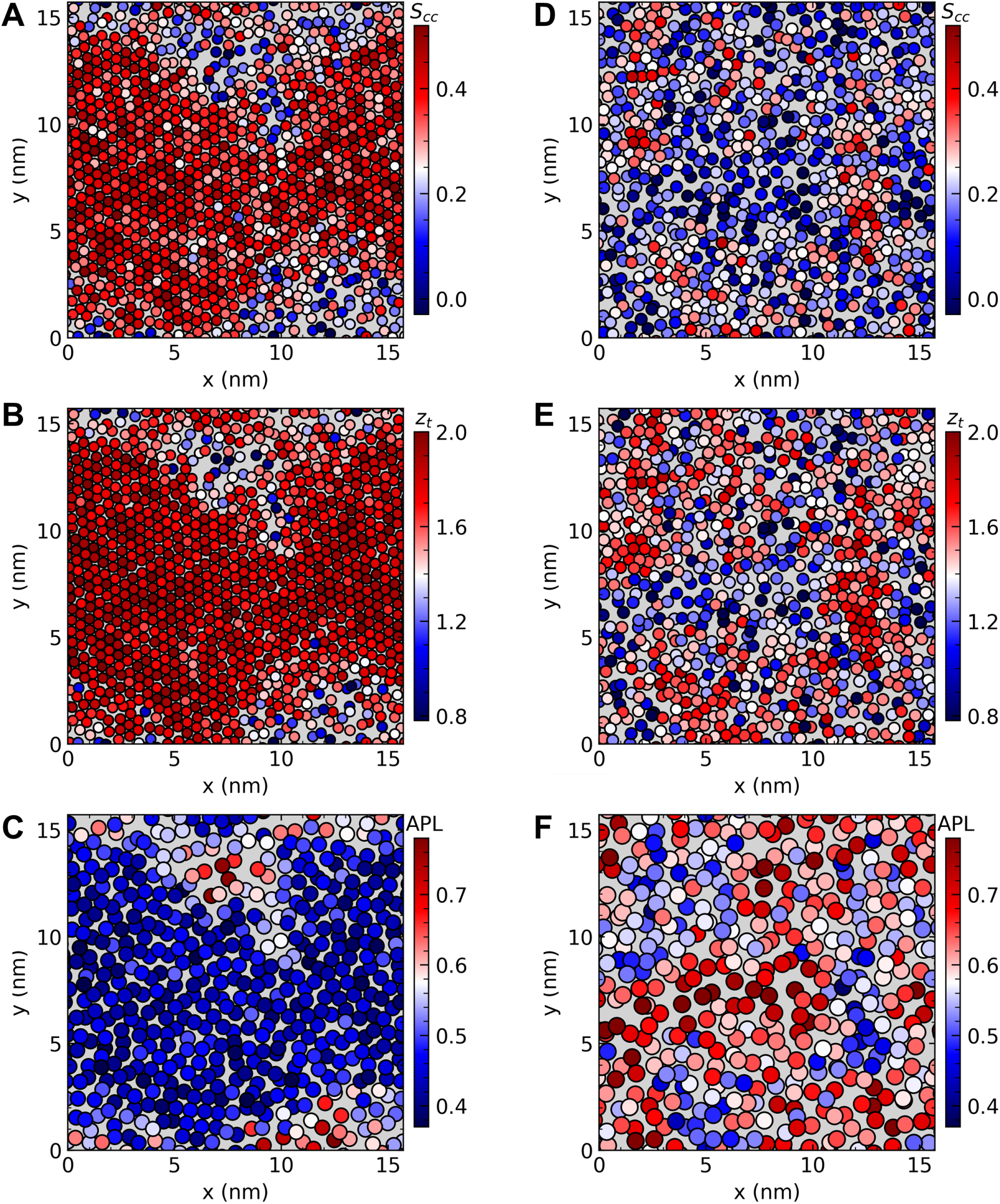
Structural comparison between the compressed and extended leaflets under stress asymmetry. Heatmaps of structural properties for a POPE membrane under number asymmetry of Δ𝑛 ≈ 13.0%, plotted for the final simulation snapshot. Panels (A-C) correspond to the compressed leaflet, and panels (D-F) to the extended leaflet. Shown are (A,D) carbon–carbon order parameter, (B,E) tail height in the z-direction, and (C,F) area per lipid. Each bead represents (A,B,D,E) the center of mass of an individual lipid tail or (C,F) the average center of mass of both tails.

**Figure S5.**
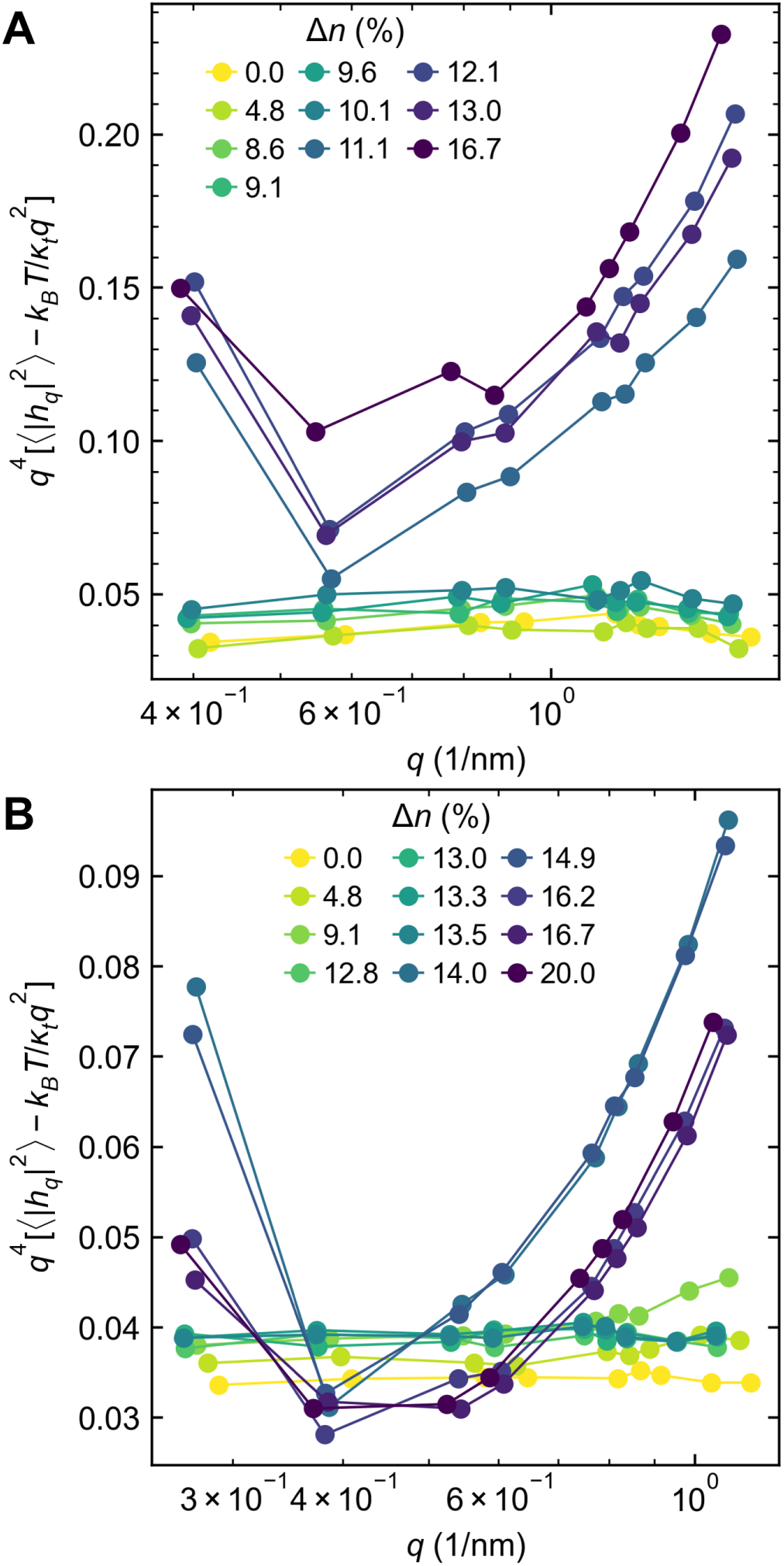
Membrane undulation spectra. Power spectra of height fluctuations for (A) all-atom POPE membranes (∼400 lipids/leaflet) and (B) MARTINI DLPC membranes (∼800 lipids/leaflet).

**Figure S6.**
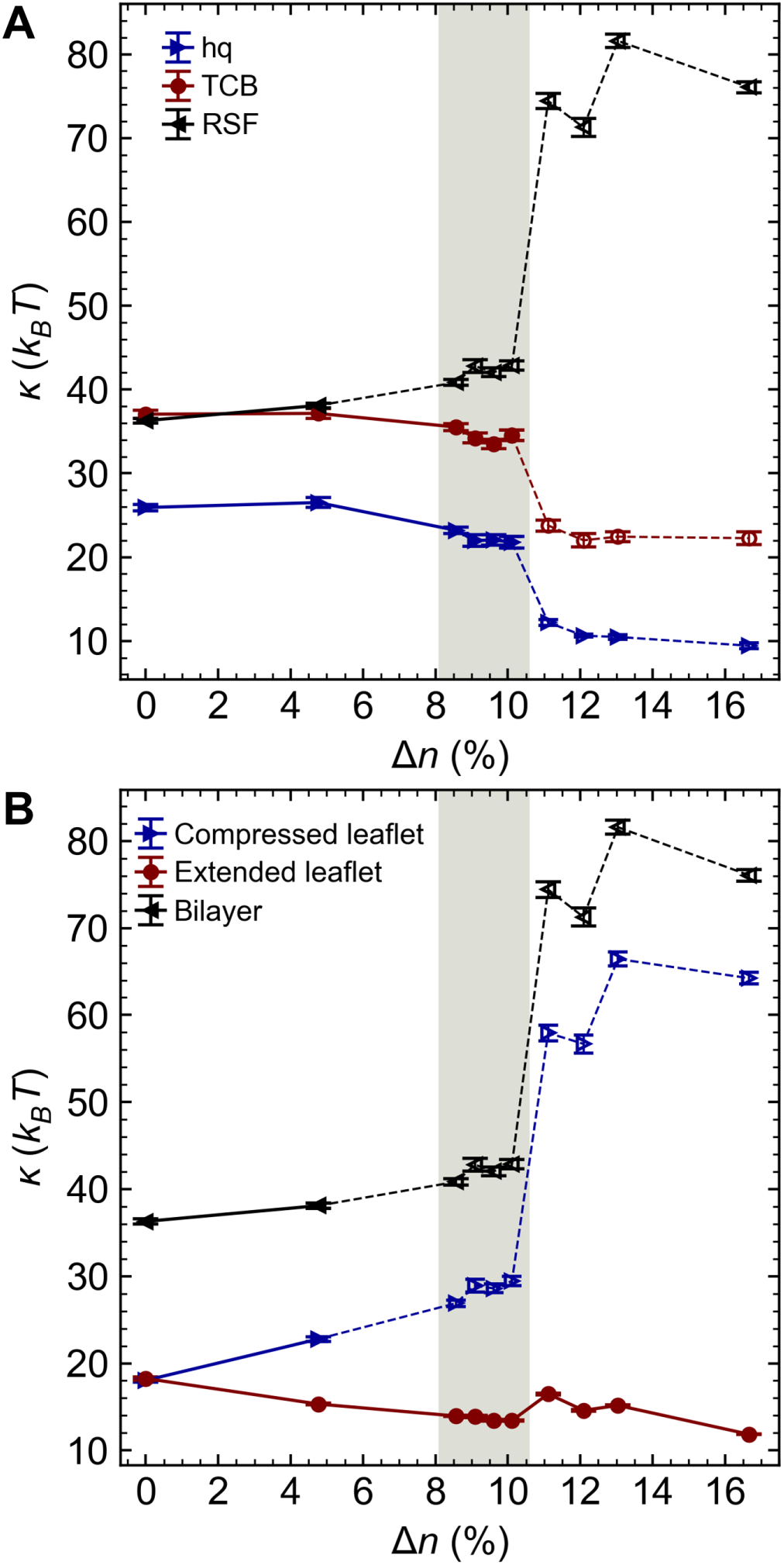
Bending moduli of all-atom POPE membranes (∼400 lipids/leaflet). (A) Comparison of bending moduli estimated using different methods as a function of number asymmetry. (B) Per-leaflet bending moduli obtained from the RSF method.

**Figure S7.**
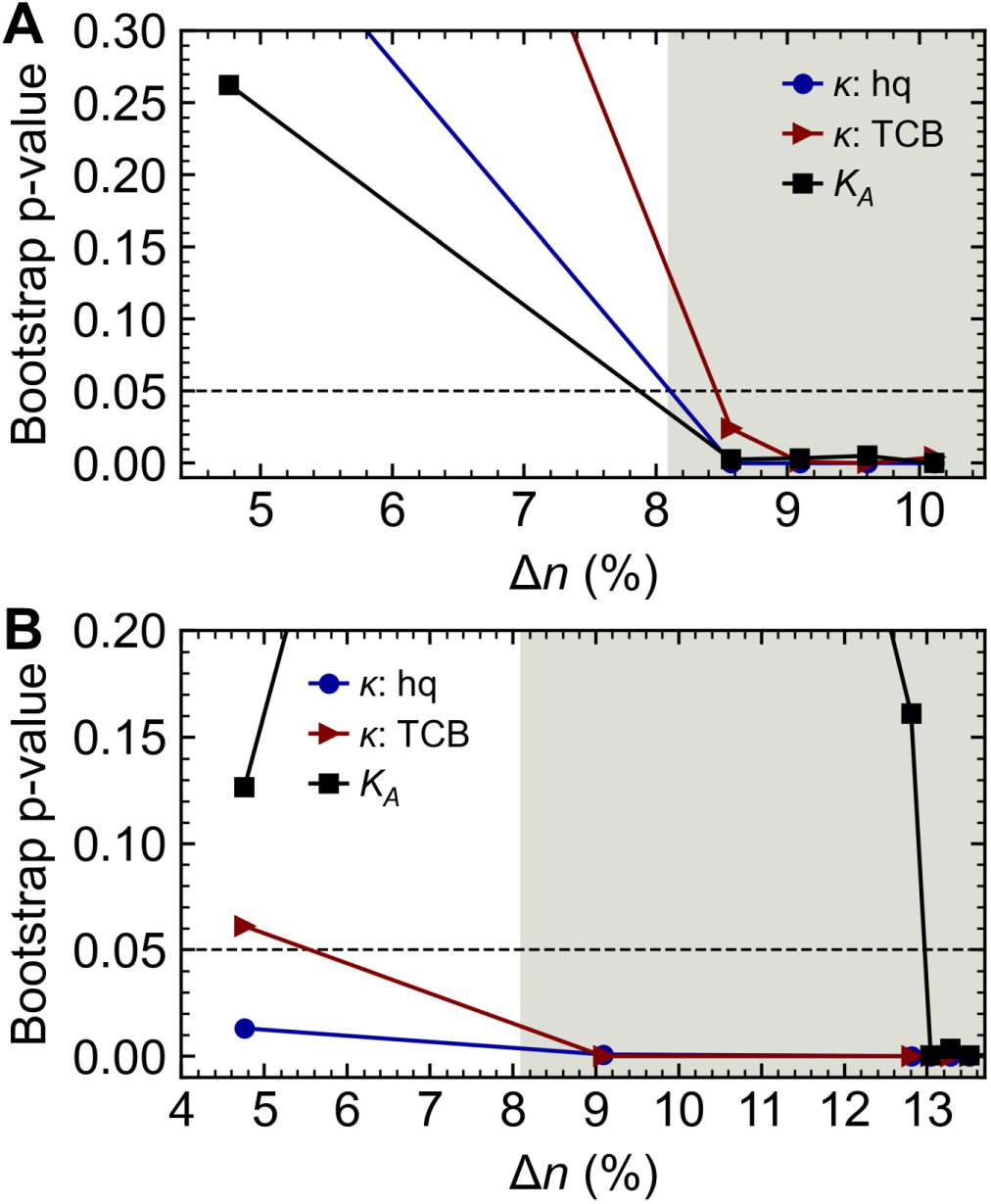
Bootstrap *p*-values for membrane elastic moduli. Results shown for (A) all-atom POPE membranes (∼400 lipids/leaflet) and (B) MARTINI DLPC membranes (∼800 lipids/leaflet).

**Figure S8.**
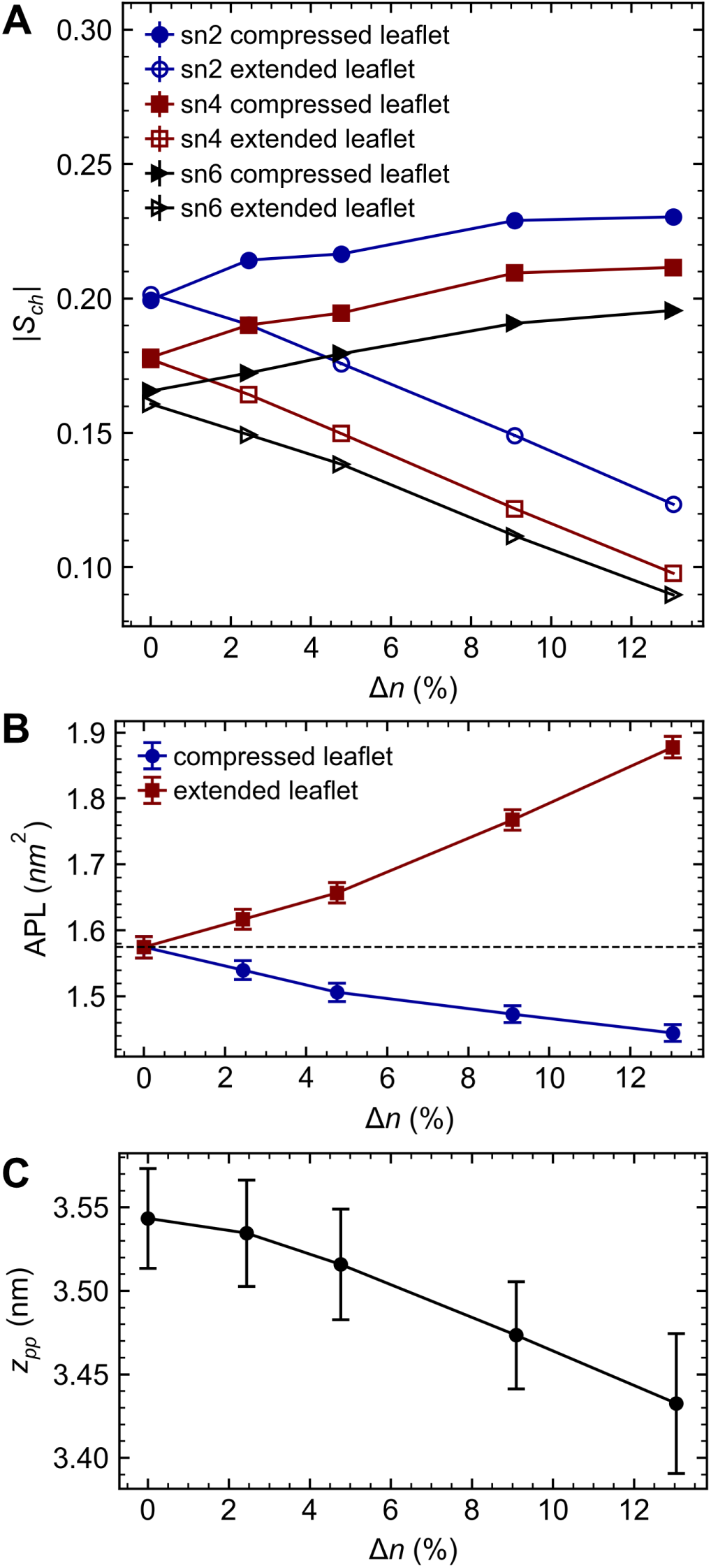
Asymmetry-induced structural changes in single-component Lipid A bilayers (310 K). (A) Carbon-hydrogen order parameter, (B) area per lipid, and (C) membrane thickness as functions of number asymmetry.

**Figure S9.**
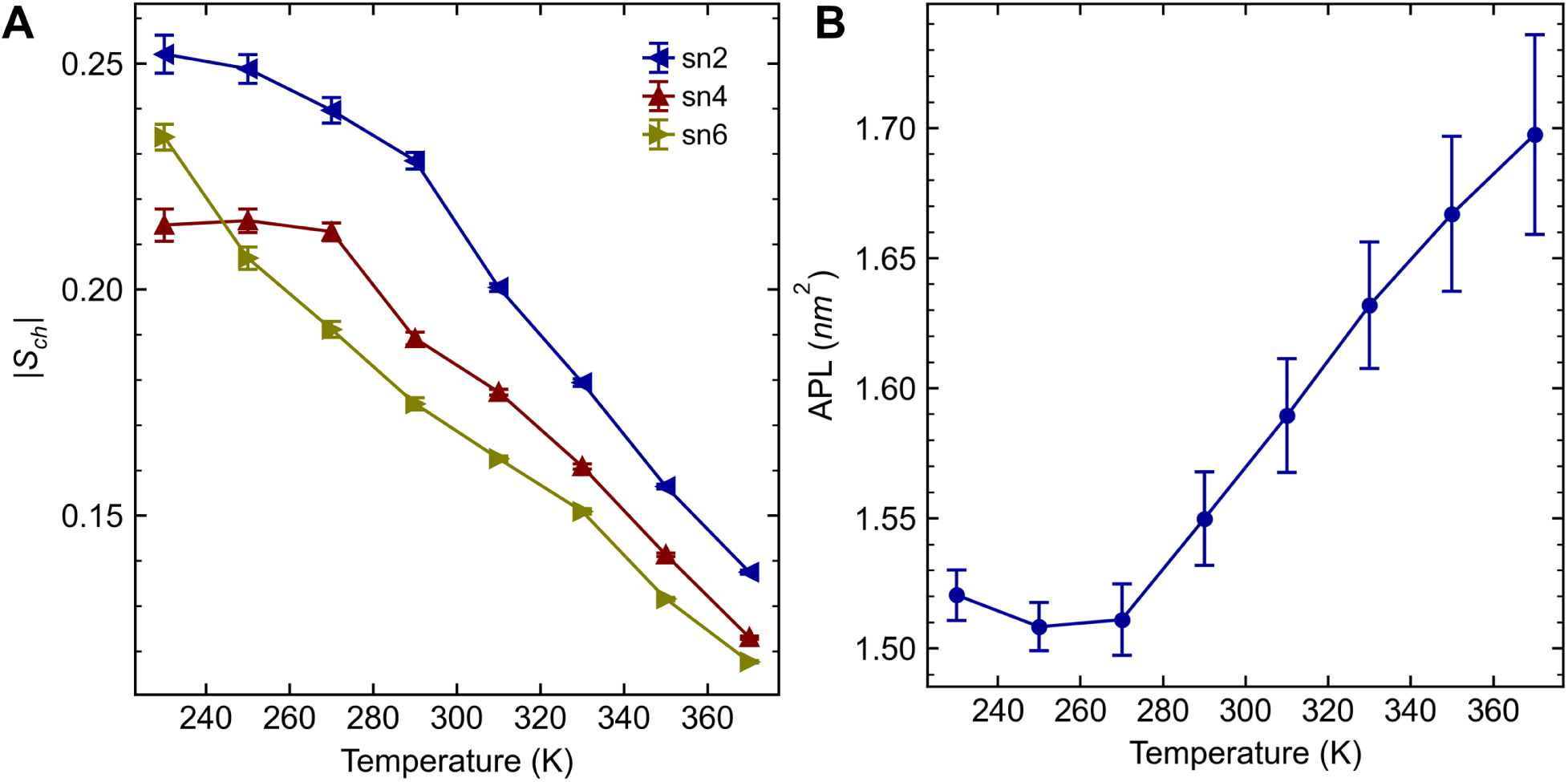
Phase transition temperature of symmetric Lipid A bilayers. (A) Carbon-hydrogen order parameter and (B) area per lipid as functions of temperature.

**Figure S10.**
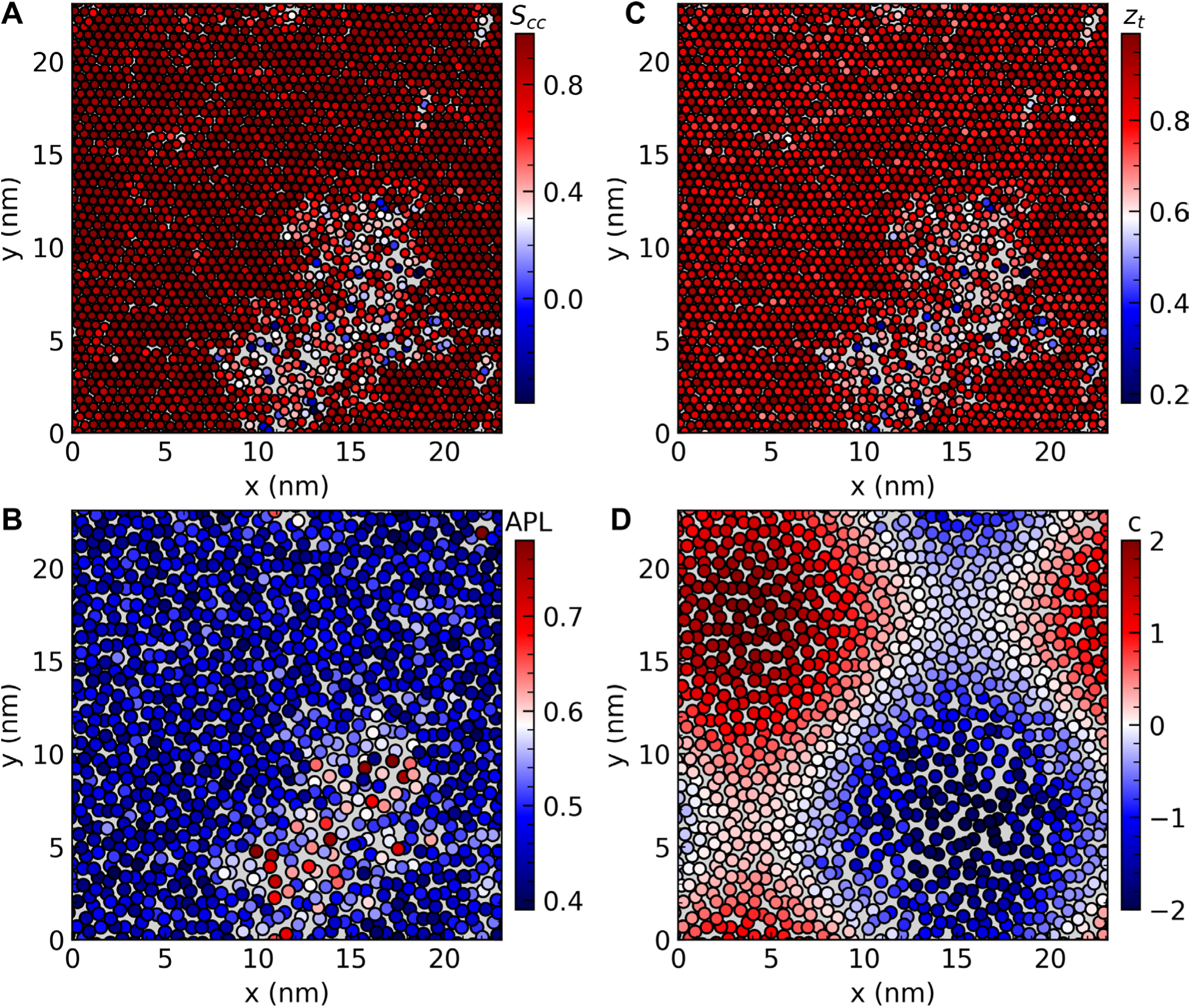
Lateral heterogeneity in MARTINI DLPC membranes under high asymmetry. Heatmaps of structural properties for a membrane with Δ𝑛 ≈ 16.7%, plotted for the final simulation snapshot: (A) carbon-carbon order parameter, (B) area per lipid, (C) tail height in the z-direction, and (D) per-lipid curvature. Each bead corresponds to (A,C) the center of mass of an individual lipid tail or (B,D) the average center of mass of both tails.

**Figure S11.**
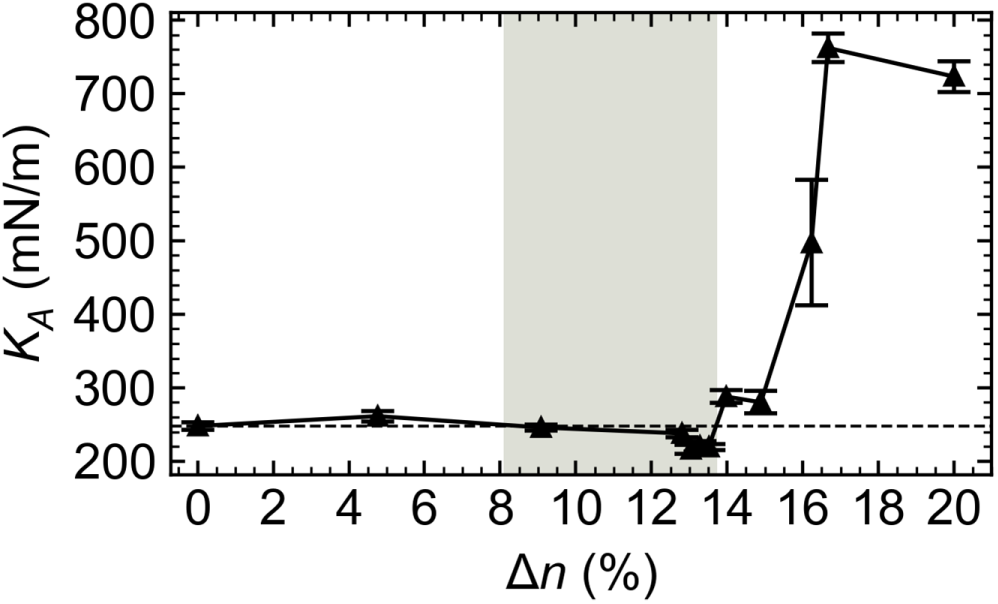
Area compressibility moduli of MARTINI DLPC membranes (∼800 lipids/leaflet). Error bars represent standard errors of the mean obtained via block averaging. Shaded regions indicate the near-transition regime characterized by transient gel-like domains.

**Figure S12.**
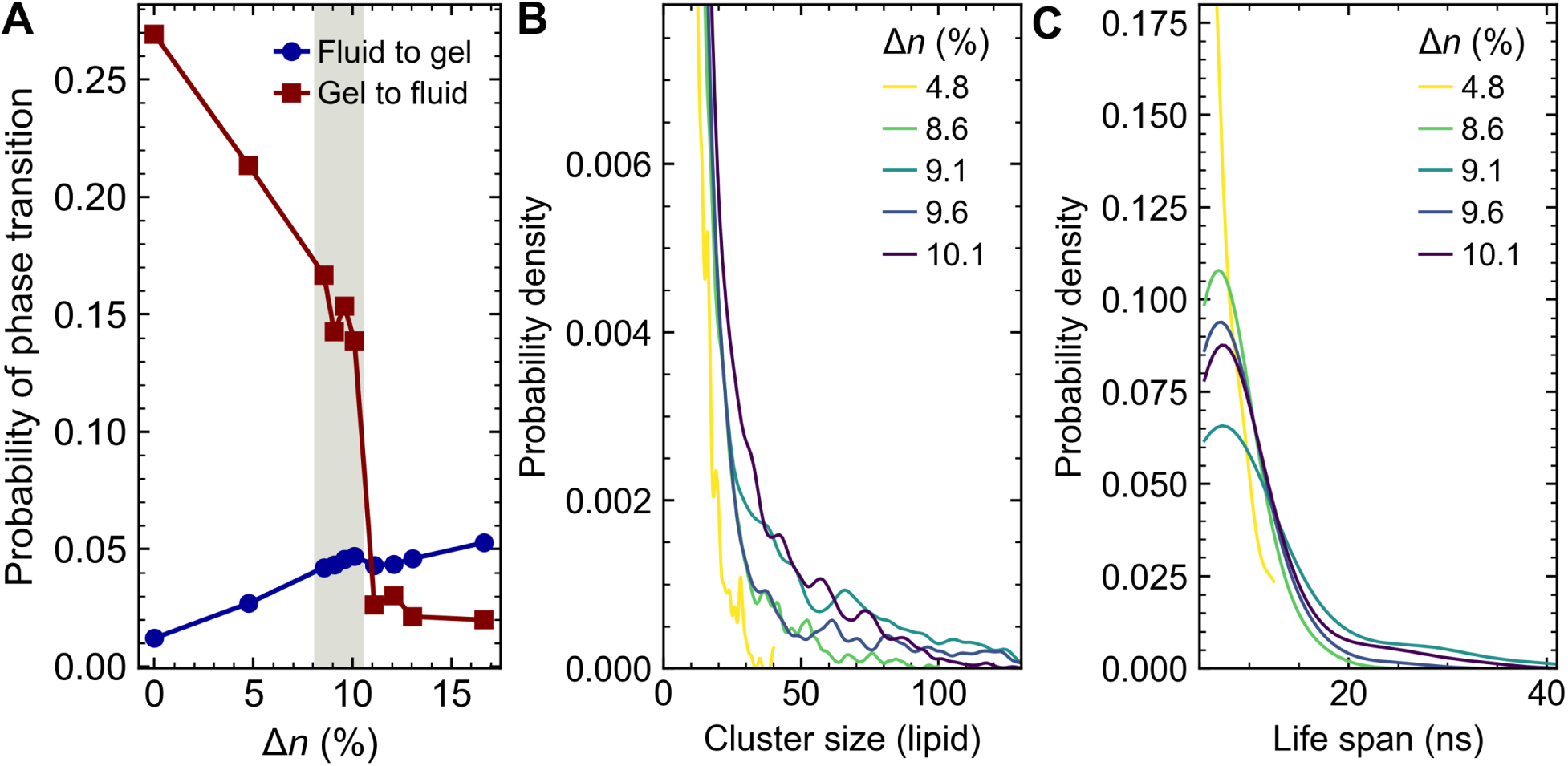
Phase statistics for all-atom POPE membranes (∼400 lipids/leaflet). (A) Probability of lipid phase switching between fluid and gel states. Probability densities of (B) gel-domain lifetimes and (C) gel-domain sizes across different number asymmetries. Shaded regions indicate the near-transition regime characterized by transient gel domains.

**Figure S13.**
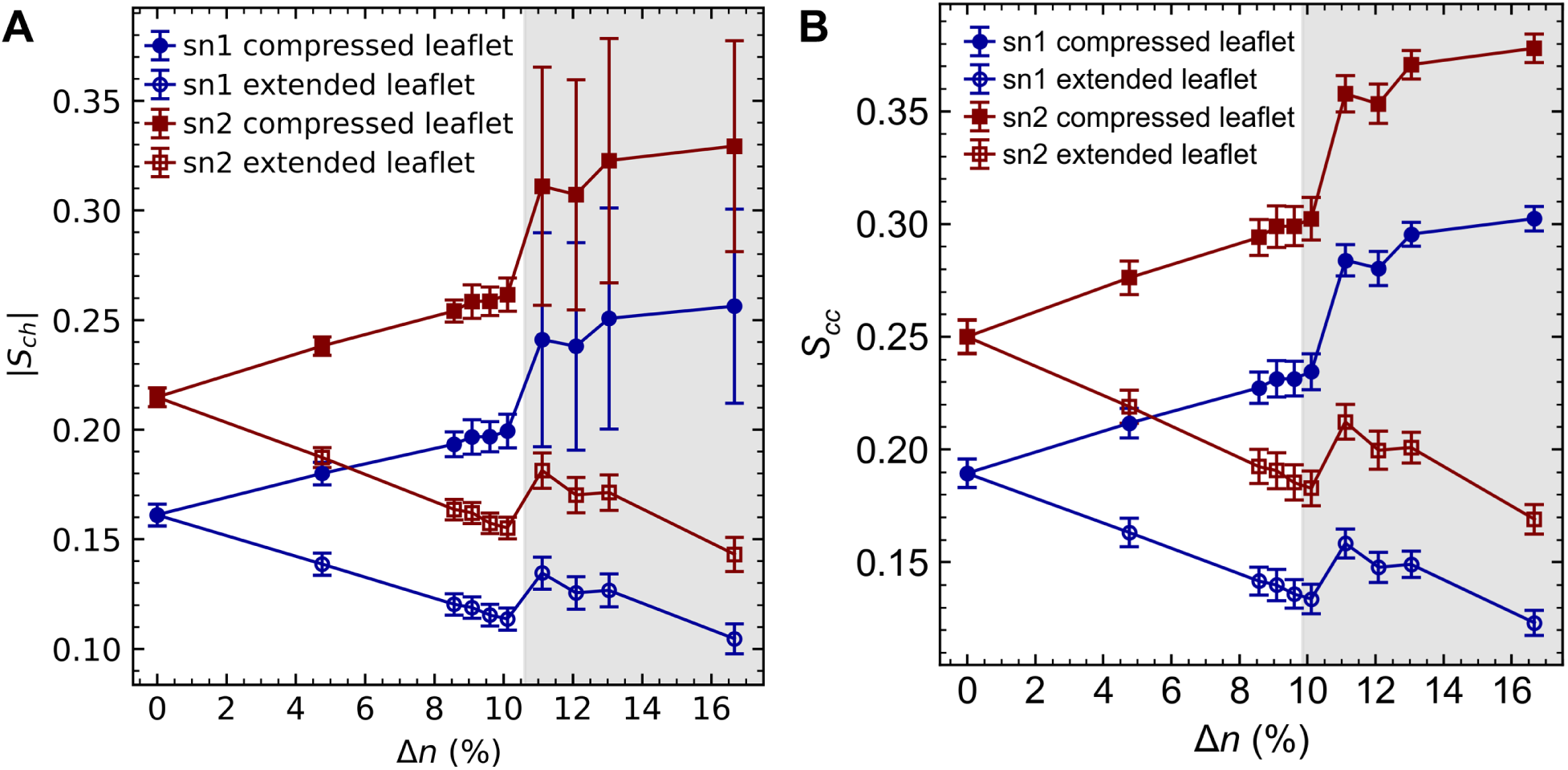
Comparison of different order parameters in POPE membranes (∼400 lipids/leaflet). (A) Carbon-hydrogen order parameter and (B) carbon-carbon order parameter as functions of number asymmetry.

**Figure S14.**
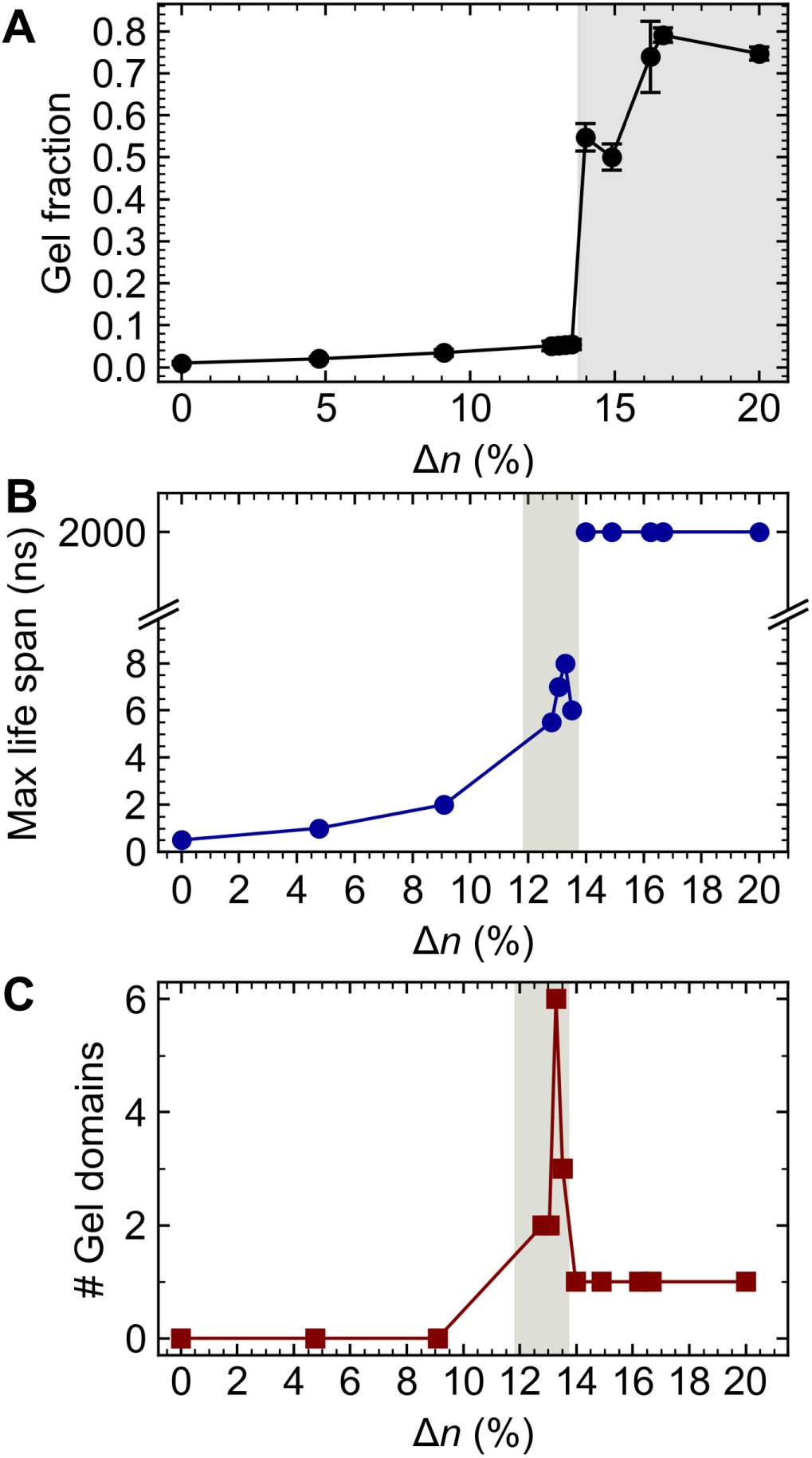
Phase properties of MARTINI DLPC membranes (∼800 lipids/leaflet). (A) Fraction of gel-phase lipids in the compressed leaflet, (B) maximal gel-cluster lifetime over a 2 μs simulation, and (C) total number of gel domains formed, all plotted as functions of number asymmetry.

## Supplemental Tables

**Table S1.**
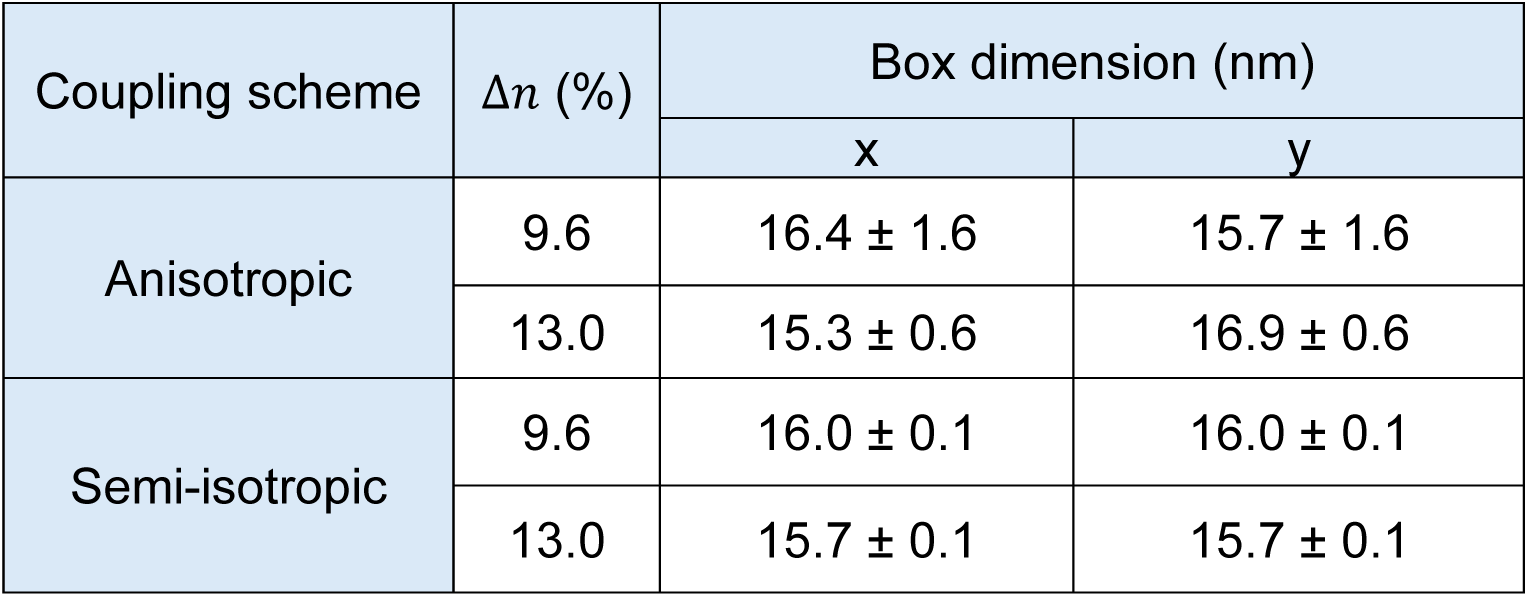
Box dimensions for selected POPE all-atom membranes under different pressure coupling schemes. Error bars indicate the standard deviations over 2 µs of sampling time.

| Coupling scheme | $\Delta n$ (%) | Box dimension (nm) | |
| --- | --- | --- | --- |
|  |  | x | y |
| Anisotropic | 9.6 | $16.4 \pm 1.6$ | $15.7 \pm 1.6$ |
| | 13.0 | $15.3 \pm 0.6$ | $16.9 \pm 0.6$ |
| Semi-isotropic | 9.6 | $16.0 \pm 0.1$ | $16.0 \pm 0.1$ |
| | 13.0 | $15.7 \pm 0.1$ | $15.7 \pm 0.1$ |

**Table S2.** Relationship between number asymmetry and differential stress for all-atom POPE membranes.

| $\Delta n$ (%) | 0 | 3.8 | 5.7 | 8.3 | 10.7 |
| --- | --- | --- | --- | --- | --- |
| $\Delta\gamma$<br>(mN/m) | 0* | -17.1 | -23.2 | -36.1 | -57.7 |

\* For symmetric systems, the differential stress was not explicitly calculated and was assumed to be zero.

**Table S3.**
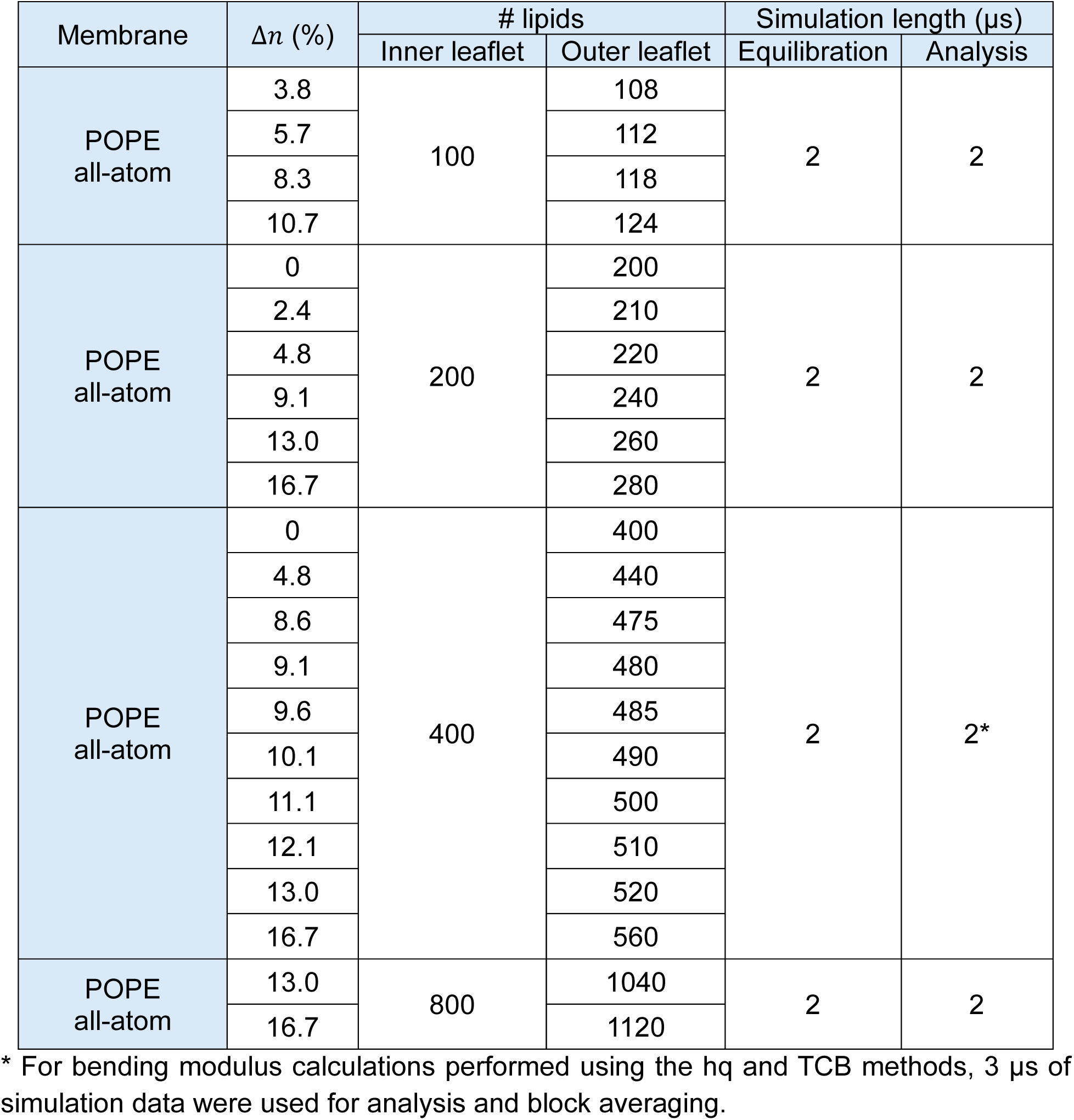
Simulation details for all POPE single-component membrane systems.

**Table S4.** Simulation details for all DLPC single-component membrane systems.

| Membrane | $\Delta n$ (%) | # lipids | | Simulation length (μs) | |
| --- | --- | --- | --- | --- | --- |
|  |  | Inner leaflet | Outer leaflet | Equilibration | Analysis |
| DLPC<br>MARTINI 2.0 | 0 | 800 | 800 | 2 | 2 |
|  | 4.7 |  | 880 |  |  |
|  | 9.1 |  | 960 |  |  |
|  | 12.8 |  | 1035 |  |  |
|  | 13.0 |  | 1040 |  |  |
|  | 13.3 |  | 1045 |  |  |
|  | 13.5 |  | 1050 |  |  |
|  | 14.0 |  | 1060 |  |  |
|  | 14.9 |  | 1080 |  |  |
|  | 16.2 |  | 1110 |  |  |
|  | 16.7 |  | 1120 |  |  |
|  | 20.0 |  | 1200 |  |  |

**Table S5.** Simulation details for all lipid A single-component membrane systems.

| Membrane | $\Delta n$ (%) | # lipids | | Simulation length (μs) | |
| --- | --- | --- | --- | --- | --- |
|  |  | Inner leaflet | Outer leaflet | Equilibration | Analysis |
| Lipid A<br>all-atom<br>($T_m$ analysis) | 0 | 36 | 36 | 5 | 1 |
| Lipid A<br>all-atom<br>(asymmetry) | 0 | 80 | 80 | 5 | 2 |
|  | 2.4 |  | 84 |  |  |
|  | 4.8 |  | 88 |  |  |
|  | 9.1 |  | 96 |  |  |
|  | 13.0 |  | 104 |  |  |
|  | 16.7 |  | 112 |  |  |

**Table S6.** Simulation details for all OM multicomponent membrane systems.

| Membrane | $\Delta n_r$ (%) | # lipids | | | Simulation length (μs) | |
| --- | --- | --- | --- | --- | --- | --- |
|  |  | Inner leaflet |  | Outer leaflet (Lipid A) | Equilibration | Analysis |
|  |  | POPE | POPG |  |  |  |
| OM all-atom | -30.0 | 283 | 128 | 216 | 5 | 2 |
|  | -25.0 | 292 | 132 | 208 |  |  |
|  | 0 | 305 | 138 | 163 |  |  |
|  | 10.0 | 358 | 162 | 174 |  |  |
|  | 20.1 | 382 | 173 | 170 |  |  |
|  | 25.0 | 365 | 165 | 156 |  |  |
|  | 30.0 | 360 | 163 | 148 |  |  |
|  | 35.1 | 369 | 167 | 146 |  |  |
| OM all-atom | -5.0 | 64 | 29 | 36 | 5 | 2 |
|  | -2.3 | 64 | 29 | 35 |  |  |
|  | 0 | 73 | 33 | 39 |  |  |
|  | 2.5 | 71 | 32 | 37 |  |  |
|  | 5.4 | 71 | 32 | 36 |  |  |

**Table S7.** Relationship between relative number asymmetry and differential stress for all-atom OM models.

|  |  |  |  |  |  |
| --- | --- | --- | --- | --- | --- |
| $\Delta n_r$ (%) | -5.0 | -2.5 | 0 | 2.3 | 5.4 |
| $\Delta\gamma$ (mN/m) | -9.5 | -3.4 | -0.4 | 5.8 | 10.2 |

## Supplemental Videos

**Movie S1.** HMM-resolved phase state and heatmaps of structural properties of each lipid in one leaflet of a symmetric all-atom POPE membrane (Δ𝑛 = 0%), visualized over 2 µs of simulation snapshots.

**Movie S2.** HMM-resolved phase state and heatmaps of structural properties of each lipid in the compressed leaflet of an all-atom POPE membrane with Δ𝑛 = 10.1%, visualized over 2 µs of simulation snapshots.

**Movie S3.** HMM-resolved phase state and heatmaps of structural properties of each lipid in the compressed leaflet of an all-atom POPE membrane with Δ𝑛 = 13.0%, visualized over 2 µs of simulation snapshots.

**Movie S4.** Head-group identity and heatmaps of structural properties of each lipid in the inner (PLs) leaflet of an all-atom outer membrane (OM) with relative number asymmetry Δ𝑛_𝑟_ = 30.0%, visualized over 2 µs of simulation snapshots.

